# Reconstructing tumor evolutionary histories and clone trees in polynomial-time with SubMARine

**DOI:** 10.1101/2020.06.11.146100

**Authors:** Linda K. Sundermann, Jeff Wintersinger, Gunnar Rätsch, Jens Stoye, Quaid Morris

## Abstract

Tumors contain multiple subpopulations of genetically distinct cancer cells. Reconstructing their evolutionary history can improve our understanding of how cancers develop and respond to treatment. Subclonal reconstruction methods cluster mutations into groups that co-occur within the same subpopulations, estimate the frequency of cells belonging to each subpopulation, and infer the ancestral relationships among the subpopulations by constructing a clone tree. However, often multiple clone trees are consistent with the data and current methods do not efficiently capture this uncertainty; nor can these methods scale to clone trees with a large number of subclonal populations.

Here, we formalize the notion of a partial clone tree that defines a subset of the pairwise ancestral relationships in a clone tree, thereby implicitly representing the set of all clone trees that have these defined pairwise relationships. Also, we introduce a special partial clone tree, the *Maximally-Constrained Ancestral Reconstruction* (MAR), which summarizes all clone trees fitting the input data equally well. Finally, we extend commonly used clone tree validity conditions to apply to partial clone trees and describe SubMARine, a polynomial-time algorithm producing the *subMAR*, which approximates the MAR and guarantees that its defined relationships are a subset of those present in the MAR. We also extend SubMARine to work with subclonal copy number aberrations and define equivalence constraints for this purpose. In contrast with other clone tree reconstruction methods, SubMARine runs in time and space that scales polynomially in the number of subclones.

We show through extensive simulation and a large lung cancer dataset that the subMAR equals the MAR in > 99.9% of cases where only a single clone tree exists and that it is a perfect match to the MAR in most of the other cases. Notably, SubMARine runs in less than 70 seconds on a single thread with less than one Gb of memory on all datasets presented in this paper, including ones with 50 nodes in a clone tree.

The freely-available open-source code implementing SubMARine can be downloaded at https://github.com/morrislab/submarine.

**Author summary:** Cancer cells accumulate mutations over time and consist of genetically distinct subpopulations. Their evolutionary history (as represented by tumor phylogenies) can be inferred from bulk cancer genome sequencing data. Current tumor phylogeny reconstruction methods have two main issues: they are slow, and they do not efficiently represent uncertainty in the reconstruction.

To address these issues, we developed SubMARine, a fast algorithm that summarizes all valid phylogenies in an intuitive format. SubMARine solved all reconstruction problems in this manuscript in less than 70 seconds, orders of magnitude faster than other methods. These reconstruction problems included those with up to 50 subclones; problems that are too large for other algorithms to even attempt. SubMARine achieves these result because, unlike other algorithms, it performs its reconstruction by identifying an upper-bound on the solution set of trees. In the vast majority of cases, this upper bound is tight: when only a single solution exists, SubMARine converges to it > 99.9% of the time; when multiple solutions exist, our algorithm correctly recovers the uncertain relationships in more than 80% of cases.

In addition to solving these two major challenges, we introduce some useful new concepts for and open research problems in the field of tumor phylogeny reconstruction. Specifically, we formalize the concept of a partial clone tree which provides a set of constraints on the solution set of clone trees; and provide a complete set of conditions under which a partial clone tree is valid. These conditions guarantee that all trees in the solution set satisfy the constraints implied by the partial clone tree.

## 1 Introduction

Tumors contain multiple, major subpopulations of genetically distinct cancer cells [1, 2]. The evolutionary history of a cancer can be reconstructed using the allelic frequencies of the clonal and subclonal mutations in one or more bulk samples of a single cancer. Multiple samples from the same individual’s cancer can be either spatially distinct [3] or longitudinal [4, 5]. *Clonal* mutations are present in all profiled cancer cells and were inherited from their most recent common ancestor; *subclonal* mutations are those that are present only in some, or one, of the subpopulations. Subclonal reconstruction algorithms infer the ancestral relationships among the subpopulations by constructing a *clone tree*; the genotypes of individual subpopulations can then be determined using this tree. These trees contribute to a better understanding of cancer development and response to treatment [6, 7] by helping to identify key steps in cancer progression [8, 9].

Clone trees are directed, rooted trees whose nodes correspond to different subclones, where directed edges link parental subclones to their direct descendants. A *subclone* is a group of cells descended from a single founder cell; and corresponds to a subtree (or clade) of the phylogeny of the cancerous subpopulations. Methods to construct clone trees assume that these cells all inherit the mutations present in the founder cells unless those mutations are removed from the cell through a copy number loss of its genomic locus. Subclones are associated with a set of subclone-defining mutations which are present in this founder cell but not in its parental subclone. The root of the tree, called the *germline*, represents the embryonic cell, which is the founder of all cancer cells (and all other cells in the body). In most, but not all cancers, there is a single cancerous subclone that is the ancestor of all the others; this special subclone is called the *clonal population* and it is associated with the cancer’s clonal mutations.

Although there has been substantial progress in developing algorithms to build clone trees from bulk tumor samples [10–22]; two key challenges remain: scaling algorithms to clone trees with many subclones and efficiently capturing uncertainty in the clone trees. These challenges persist even when mutation allele frequency measurements are very precise. Here we address these two challenges: first assuming perfect accuracy in the allele frequencies and second exploring relaxing that assumption by introducing a noise buffer. Specifically, we introduce an algorithm, SubMARine, which runs in polynomial-time and summarizes an upper bound on the solution set of clone trees for an input set of subclonal frequencies using a partial clone tree, a new data structure that defines the ancestral relationships between the pairs of subclones.

### Contributions

Here we introduce and formalize the notion of a partially-defined clone tree, or *partial clone tree* for short. This representation is a partial solution to a clone tree reconstruction problem that defines a subset of the pairwise ancestral relationships between the subclones, as well as a set of potential parents for each subclone. A partial clone tree is not a tree itself, but it implicitly defines a set of clone trees, i. e., all those trees that (i) are consistent with the ancestral relationships defined in the partial clone tree and (ii) select their parents from the possible parent set. The partial clone tree is thus a polynomial-space representation of a potentially exponentially-sized set of clone trees.

We also introduce a special partial clone tree: the *Maximally-Constrained Ancestral Reconstruction*, or MAR for short, which provides a complete summary of pairwise ancestral relationship constrained by the input data. Specifically, when multiple clone trees provide identically good fits to the mutation allele frequency data, the MAR captures all (and only) the pairwise ancestral relationships shared by this solution set of clone trees.

Additionally, we describe a polynomial-time algorithm, SubMARine, that produces the *subMAR*, which approximates the MAR. The ancestral relationships defined in the subMAR are guaranteed to be subset of those present in the MAR. Through extensive simulation and in a large real dataset, we demonstrate that the subMAR almost always perfectly recovers the MAR. In particular, when the MAR represents a single clone tree solution, the subMAR matches it in > 99.9% of our experiments. SubMARine is designed not only for the basic clone tree reconstruction problems commonly addressed by other approaches, but also for more complex problems that are less often considered. The basic problems include only simple somatic mutations (SSMs), including single nucleotide variants and small insertions and deletions, and clonal copy number aberrations (CNAs). The extended version of SubMARine also considers subclonal CNAs. Notably, SubMARine runs in less than 70 seconds on a single thread with less than one Gb of memory on all datasets presented in this paper, including ones with up to 50 subclones.

Finally, although SubMARine is primarily designed assuming that the input subclonal frequencies are precisely measured and hence constant, we also introduce a noise-buffered version of SubMARine. This version estimates the minimum deviation required from the input frequencies for a valid partial clone tree to exist. The noise-free version of SubMARine is immediately applicable to many real clone tree reconstruction problems without modification. In the discussion section, we discuss strategies to use noise-buffered SubMARine to explore the space of clone trees with good fits to the input frequencies.

## 2 Background

To define CNAs, the genome is divided into *segments*, with neighboring segments having different allele-specific average copy numbers in one or more samples. CNA reconstruction algorithms identify these segments and infer the average allele-specific copy numbers within them [23, 24]. However, fewer algorithms indicate the evolutionary relationship among the CNAs [10, 14, 22, 25]. SSMs are quantified experimentally by reporting their variant allele frequencies (VAFs) in each sample as estimated by short-read sequencing. These VAFs can be transformed into estimates of the cellular frequency of the SSMs by accounting for clonal CNAs in the sample influencing this transformation [26]. SSMs can be grouped into subclones based on these inferred cellular frequencies, thus estimating the associated subclonal frequencies in each sample [27–29]. With some modifications, similar algorithms can also be used to group CNAs into subclones [30–32]. The accuracy of the cellular frequency estimates, CNA reconstructions, and subclonal groupings depends heavily on the sequencing depth, degree of aneuploidy, and purity of the samples [33]. However, even under the best of conditions, when there is high accuracy in all of these, there remain substantial challenges in clone tree reconstruction.

Figure S1 shows a clone tree that solves a clone tree reconstruction problem by representing the ancestral relationships among the subclones. The solution to a **clone tree reconstruction problem** is a valid clone tree for the following input, which can be derived from a subclonal reconstruction problem: *K* subclones (including the germline); their subclonal frequencies in each of *N* samples, represented by the subclonal frequency matrix 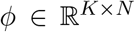; *L* CNAs assigned to segments, subclones and parental alleles; and *J* SSMs assigned to segments and subclones.

A clone tree is **valid** if it satisfies the tree, the lost allele, and the sum constraints. The **tree constraint** simply requires the clone tree, thus the ancestral relationships, to be consistent with an arborescence (i.e., a directed tree whose edges all point away from the root) whose root is the germline. The **lost allele constraint**, which applies to both CNAs and SSMs, insists that mutations cannot occur on segments lost in an ancestral cell (see Section S2 for more details). Finally, because subclones represent subtrees (or clades) of phylogenies, the subclonal frequencies of a subclone must be larger than or equal to the sum of frequencies of its children in all samples, hence a **sum constraint** [11, 15] on the frequencies must hold in the clone tree:

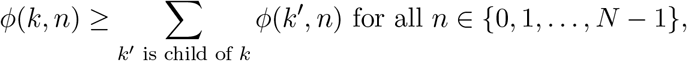

where 0 ≤ *ϕ*(*k, n*) ≤ 1 is the frequency of subclone *k* in sample *n*.

The **basic clone tree reconstruction problem** considers only SSMs and clonal CNAs and, as such, only needs to consider *ϕ* when searching for valid clone trees. This problem was shown to be NP-complete [11]. The **extended clone tree reconstruction problem**, introduced here, requires additional input, including an impact matrix 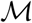. We introduce the extended problem in Section 4.

### Previous work

Often, multiple clone trees solve a clone tree reconstruction problem because the input data does not provide sufficient constraints to select a single solution [12, 15, 34]. The theoretical implications of this were first formally studied in [35, 36]. When there are multiple solutions, clone tree reconstruction algorithms invent other criteria to select a single solution [13, 21, 34] or they report a (hopefully) representative subset of the solution set [10, 14, 15, 18, 20]. Other methods simply enumerate all possible clone tree solutions [11, 12, 19]; however, because the solution space of clone trees grows exponentially with the number of subclones, these enumeration methods are limited to problems with a small number of subclones.

Given multiple clone trees as input, some methods identify a single [37] or multiple [38] representative consensus trees in order to capture topological features of the solution space. However, a single consensus tree cannot represent ambiguity in the data, and optimal selection of multiple consensus trees is NP-hard [38]. Furthermore, these methods already require the potentially exponentially-sized solution set of clone trees to be enumerated as input. In fact, already the problem of counting the number of valid solutions to the basic clone tree reconstruction problem is #P-complete [36].

## 3 Partial clone trees

A *partial clone tree* defines some but, generally, not all of the pairwise ancestral relationships between subclones. A defined relationship either requires one of the subclones to be an ancestor of the other, or requires that the subclone *not* be an ancestor of the other. Thus, a partial clone tree can be represented with an *ancestry matrix Z* ∈ {1, 0, −1}^*K×K*^, where:

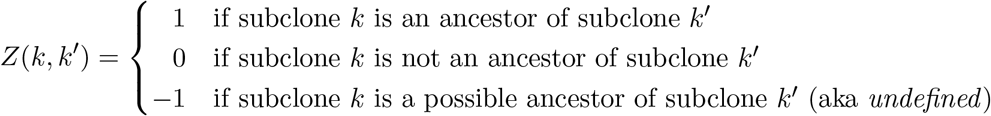

A (full) clone tree *completes* a partial clone tree if its implied pairwise ancestral relationships are consistent with the defined (i. e. non-negative) entries in *Z*. A partial clone tree thus implicitly represents the set of clone trees that complete it. Hence, a partial clone tree can be used to solve the *Maximally-Constrained Ancestral Reconstruction Problem*:

#### Problem 1 (Basic maximally-constrained ancestral reconstruction problem).

Given the subclonal frequency matrix *ϕ* of a basic clone tree reconstruction problem *t*, identify the pairwise ancestral relationships between subclones present in all valid clone trees.

The **basic maximally-constrained ancestral reconstruction (MAR)** is the unique partial clone tree that solves this problem by defining the maximal set of all of the ancestral relationships shared by the solution set of clone trees for *t*, and leaving undefined all relationships that vary within the solution set (see Figure 1). Note, however, that this does not necessarily mean that all clone trees that complete the MAR are solutions of *t*; but often they are (see Figure S2). Note also that the partial clone trees produced by SubMARine also include a possible parent matrix *τ*, which further constrains the space of completing clone trees (see Sections 3.2 and S4.2 for more details); however, this matrix is not required in the definition of the MAR.

**Fig. 1.**
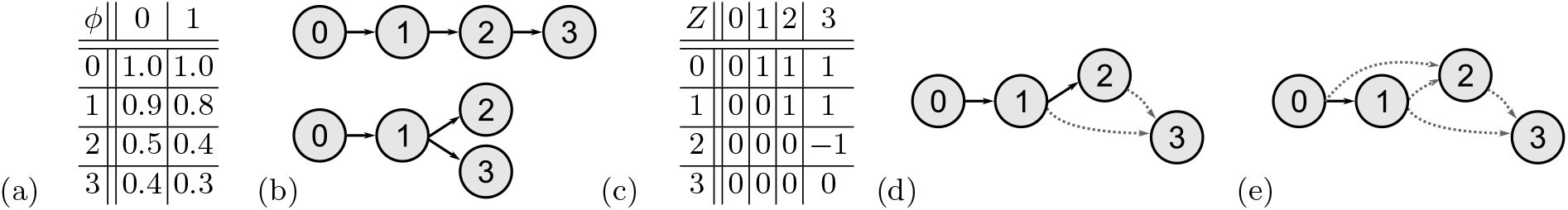
Example of a MAR for a basic maximally-constrained ancestral reconstruction problem. (a) The subclonal frequency matrix *ϕ* for the germline with index 0 and three subclones with indices 1-3 with their frequencies in two samples. (b) Set of valid clone trees that fit *ϕ*. (c) The MAR summarizing the two clone trees, represented as ancestry matrix *Z*. Whenever subclone *k* is an ancestor of subclone *k′* in both clone trees of (b), *Z*(*k,k′*) = 1. If *k* is not an ancestor of *k′* in both clones trees, *Z*(*k,k′*) = 0. If *k* is an ancestor of *k′* in one clone tree but not in the other, as for subclones 2 and 3, *Z*(*k,k′*) = −1. (d) The MAR drawn as a partial clone tree. Solid edges connect parents to their definite children (see Equation 2), dashed edges connect possible parents to their possible children (see Definition 1). (e) A partial clone tree that does not equal the MAR. Here, subclone 1 is only a possible ancestor of subclone 2, although subclone 1 is the definite ancestor in both clone trees in (b). Hence, the defined set of ancestral relationships is not maximal.

Partial clone trees generalize ancestry graphs (or evolutionary constraint networks) used by previous algorithms [11, 12, 19] as a starting point for enumerating all valid clone trees. An ancestry graph is a directed, acyclic graph (DAG), in which two subclones *k* and *k*′ are connected by an edge if *k* is a possible parent of *k*′. In these graphs, *k* is a possible parent of *k*′ if there exists no sample *n* such that *ϕ*(*k, n*) < *ϕ*(*k′, n*) (applying one aspect of the sum constraint) and if *k′* does not contain any mutation that is already lost in *k*. Clone trees can be enumerated as spanning trees with a Gabow-Myers-based algorithm [39]; they are valid if the sum constraint is satisfied for each subclone and all its children. Ancestry graphs can be represented by a partial clone tree where *Z*(*k,k′*) = −1 whenever an edge connects *k* to *k′*, and where *Z*(*k,k′*) = 0 otherwise. However, the semantics of a partial clone tree, which represents constraints on the ancestry, are not the same as the ones of an ancestry graph, which connects children to possible parents. Hence, not every ancestry matrix *Z* with only 0 and −1 entries corresponds to an ancestry graph. Also, when a partial clone tree is represented as a DAG, not every spanning tree satisfying the sum constraint completes *Z* (see Section S3.1). Here, we extend this earlier work to include ancestry relationships that must be present (i.e., *Z*(*k,k′*) = 1). Doing so allows us to not only more highly constrain the space of clone trees but also to propagate an initial set of defined ancestral relationships in *Z* to infer other ancestral relationships that must appear in the MAR. We describe SubMARine, an algorithm that allows this propagation, in Section 3.2.

### 3.1 Applying validity constraints to partial clone trees

A key contribution of this paper is the observation that the validity constraints for clone trees can be applied to partial clone trees in order to rule out, or rule in, some pairwise ancestral relationships. In addition to the sum constraint, which is already applied in the construction of ancestry graphs, SubMARine enforces the tree constraint on *Z*. This allows to rule in certain ancestral relationships, i.e., identify pairs of subclones *k* and *k′* where *Z*(*k,k′*) = 1. Doing so permits us to define, for some subclones, a set of definite child subclones they have in every solution to a basic clone tree reconstruction problem *t*; which places further constraints on *Z*.

The tree constraint requires the clone tree to be an arborescence with the germline as the root. If we define clone 0 as representing the germline, we can immediately set *Z*(0,*k*) = 1 for *k* > 0 because the root is the ancestor of all nodes in the arborescence. This first consequence of the tree constraint is called the **germline constraint**. To simplify our presentation, we also assume that subclones 1 to *K* − 1 are sorted in decreasing order of their average subclonal frequencies across samples. As an obvious consequence of the sum constraint, this ensures that *Z*(*k,k′*) = 0 whenever *k*^1^ ≤ *k*. Another consequence of the tree constraint arises from the fact that although all arborescences correspond to a unique, fully defined ancestry matrix *Z*; not all fully defined *Z* matrices correspond to arborescences. To ensure a given *Z* does represent such a tree, i. e., that it is transitive and each node has exactly one parent, it suffices to require that all the elements in *Z* satisfy a single **partial tree constraint** (see Section S3.2 for details):

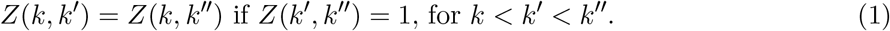

SubMARine can thus apply this constraint to partial clone trees to define an element of *Z* whenever *Z*(*k′,k″*) = 1 and either *Z*(*k,k′*) = −1 or *Z*(*k,k″*) = −1 but not both.

To assist in applying the sum constraint to partial clone trees, we define a set of *definite* children of a subclone *k*. The definite children of a subclone *k, χ*(*k*), are the set of subclones whose parent can only be *k* given the defined entries in *Z*:

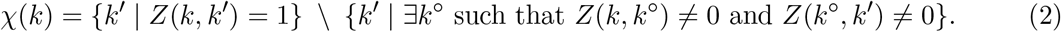

In other words, a subclone *k′* is a definite child of subclone *k* if *k* is its ancestor, and *k′* has no other (possible) ancestors that are (possible) descendants of *k*. (For Figure 1, the germline has only one definite child, which is subclone 1. Subclone 1 has subclone 2 as definite child, subclone 3 is a possible child of both subclones 1 and 2.) Thus, we can formulate the **generalized sum constraint** based simply on the set of definite children of a subclone:

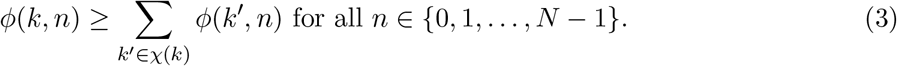

Note that when there are no undefined states in *Z, χ*(*k*) is simply the set of all children of *k*. The **lost allele constraint** can be applied without any changes to a partial clone tree (see Section S3.3).

Given these extended definitions of the validity constraints, we can now deem a partial clone tree to be *valid* if it satisfies the germline, generalized sum, lost allele, and partial tree constraints. We here note two things. First, the MAR is valid per construction (see Section S3.4). Second, when *Z* contains undefined states, some subclones have multiple possible parents and are not definite children of any subclone, hence these subclones are not considered in the generalized sum constraint. Thus, it is possible that a valid partial clone tree has no valid completions (see Figure S3).

### 3.2 SubMARine: Approximating the MAR

SubMARine is a polynomial-time algorithm that constructs the *subMAR*, which is a partial clone tree that approximates the MAR. Here we describe the basic SubMARine algorithm, which approximates the solution to the basic maximally-constrained ancestral reconstruction problem. In the following section, we describe the extended version of SubMARine.

For a basic clone tree reconstruction problem *t*, the subMAR has three important properties, which we prove in this section: it is unique, its defined ancestral relationships are a subset of those in the MAR, and as such, all valid clone trees of *t* are completions of the subMAR.

#### Algorithm 1 Functional description of the SubMARine algorithm in basic mode

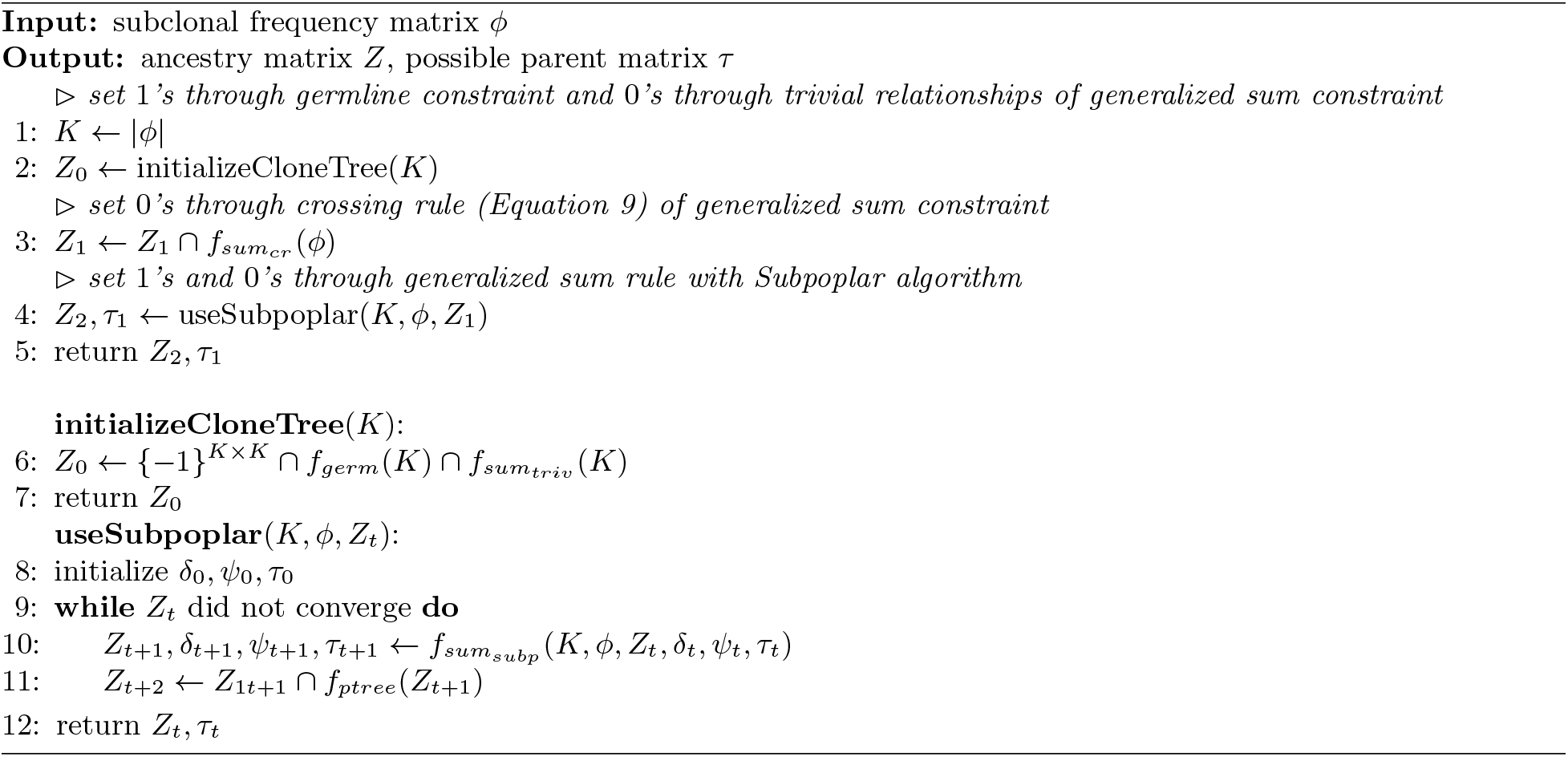

SubMARine takes as input the subclonal frequency matrix *ϕ* of a basic clone tree reconstruction problem *t*, and builds a partial clone tree by creating an ancestry matrix *Z* (see Algorithm 1 and Figure S4). Initially, this matrix contains only undefined ancestral relationships. By applying inference rules derived from the validity constraints, SubMARine updates undefined values to defined ones whenever necessary, i.e. whenever undefined values violate constraints (see Table 1). In a preprocessing phase, SubMARine applies the germline rule, setting *Z*(0,*k*) = 1 for all *k* > 0. Furthermore, all entries *Z*(*k′, k*), with *k′ ≥ k*, are set to 0 resulting from the sorting of subclones in decreasing order of their subclonal frequencies across samples and as a consequence of the generalized sum constraint. Then, the main phase of the algorithm begins by applying the crossing rule that sets *Z*(*k,k′*) = 0 for *k < k′* whenever a sample *n* exists such that *ϕ*(*k,n*) < *ϕ*(*k′,n*), as also required by the sum constraint. Afterwards, the last part of the generalized sum rule is propagated with our Subpoplar algorithm, which also propagates the partial tree constraint. This algorithm identifies definite children and rules out possible children. Its propagations lead to updates on *Z* and on the set of possible and definite parents of each subclone, which is tracked in the possible parent matrix *τ*. This tracking is necessary because the generalized sum rule can exclude possible parents for a subclone without requiring specific pairwise ancestral relationships (i. e. a subclone *k* that cannot be a possible parent of subclone *k′* can still be its possible ancestor). Whenever Subpoplar updates a relationship because of the generalized sum rule, the partial tree rule is propagated. When no more relationships can be defined through the inference rules, *Z* converged and is output as the subMAR, together with the possible parent matrix *τ*. Sections S4.1 and S4.2 provide a more detailed descriptions of SubMARine and Subpoplar, along with an analysis of their polynomial runtime.

**Table 1.**
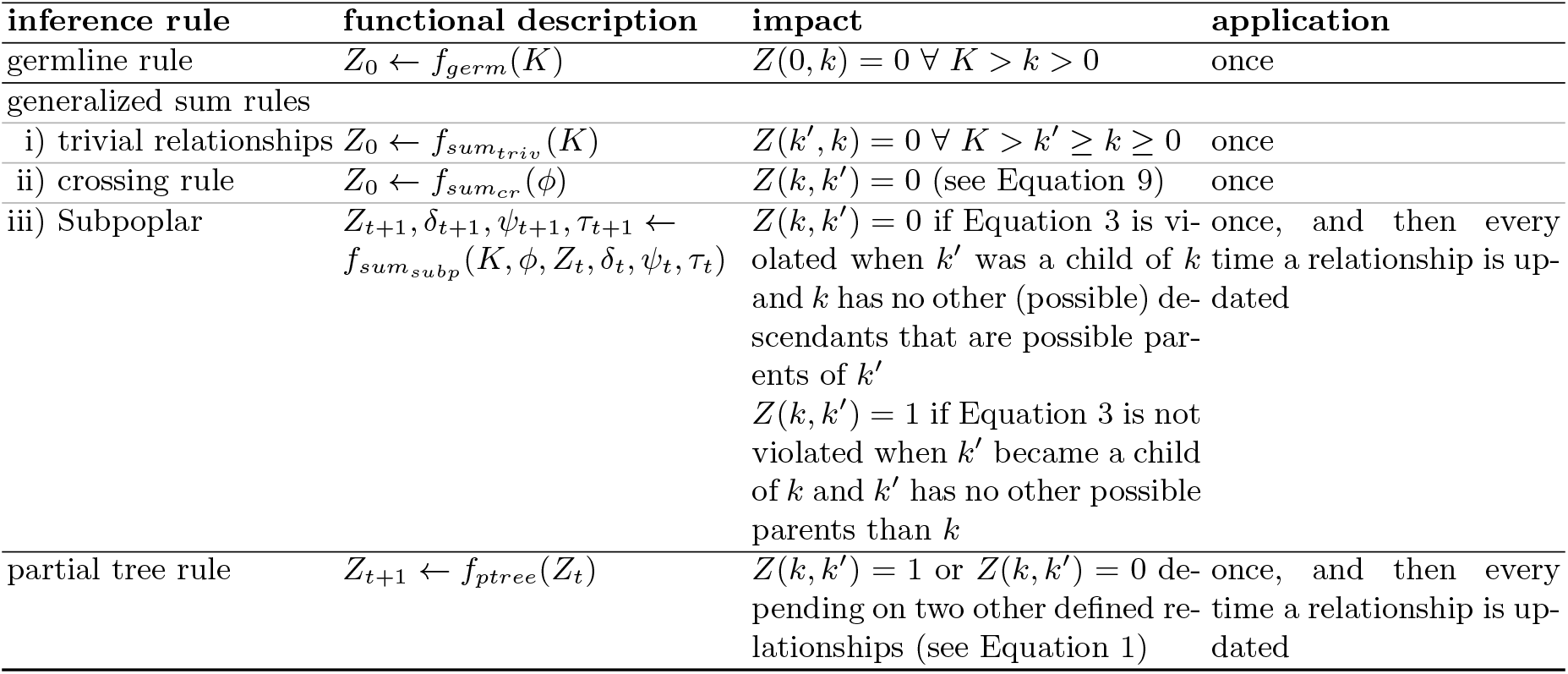
Overview of inference rules derived from the germline, generalized sum and partial tree constraint. For explanation of available frequency δ, definite parent Vector ψ and possible parent matrix *τ* see Section S4.2. *Z* is the ancestry matrix, *ϕ* the subclonal frequency matrix and *K* the number of subclones. Indices *k* and *k′* refer to subclones ordered by their average frequencies.

Note that SubMARine always converges because only undefined values are updated to defined ones and their number is finite. At convergence, *Z* represents a valid partial clone tree. If the subclonal frequency matrix *ϕ* does not support a valid partial clone tree – if, for example, one inference rule requires *Z*(*k,k′*) = 0 and another requires *Z*(*k,k′*) = 1, then SubMARine terminates and indicates the pair (*k, k′*) having a validity constraint violation. If the violation results from a generalized sum rule violation, it may be because the subclonal frequencies are not measured precisely but are actually inferred from noisy mutational frequencies. To address this issue, we describe a noise-buffered version of SubMARine in Section S4.3. In polynomial time, this version finds a minimum noise buffer that is added uniformly to parental subclonal frequencies in order to permit a valid partial clone tree. Starting from the subMAR computed with this uniform buffer, SubMARine can also find a subclone- and sample-specific noise buffer set and its corresponding subMAR, such that all completing clone trees make as little use of the buffers as possible. If the data allows, this can be done in polynomial time. Otherwise, a more exhaustive search is necessary.

If the user decides to specify additional ancestral relationships for *Z*, they are added after the preprocessing phase, followed by a propagation of the partial tree rule (see Figure S4 and Section S4.1). Furthermore, the partial tree rule is already propagated when applying the crossing rule. As additional input, clonal CNAs and SSMs can be provided. SubMARine checks then whether any SSMs are assigned to deleted segments and thus invalidate all clone trees through violating the lost allele constraint (see Section S4.4). If this is not the case, the algorithm can proceed as previously described.

#### Correctness

As described previously, the inference rules used by SubMARine change only undefined ancestral relationships to defined ones and only when, given all of the other defined relationships, one of the two possible defined ancestral relationships causes a violation of the validity constraints. So, given a starting set of defined relationships associated with *t*, each relationship defined by one of SubMARine’s inference rules is required in all valid clone trees that solve *t*. Thus, the subMAR’s defined relationships are a subset of those in the MAR.

The constructed subMAR, given the subclonal frequency matrix *ϕ* of *t*, is unique because the order in which the inference rules get applied does not matter as long as all rules are applied and propagated until convergence. It is easy to show that order of application is unimportant. Imagine a case where SubMARine generates two different subMARs, both starting from the same initial set of defined relationships, but that differ due to the order in which the inference rules were applied. Because each subMAR’s defined relationships are a subset of those in the MAR, so long as the MAR is defined (i.e., there is at least one valid and complete clone tree solution), all pairwise relationships that differ between these two subMARs are defined in one subMAR and undefined in the other. None of SubMARine’s inference rules depend on an undefined relationship in order to update another undefined relationship. As such, there must be a path of inference rules linking all defined relationships shared by the two subMARs to each defined relationship unique to one of the two subMARs. Because this path exists, and the relationship is undefined in one of the two subMARs, the inference rules have not been propagated to convergence in the subMAR where the relationship is undefined. Ergo, so long as the inference rules are propagate to convergence, and the MAR is defined, two subMARs generated from the same starting point, using the same rules, are identical. As such, the subMAR is unique.

In summary, because (i) all ancestral relationships defined in the subMAR are a subset of those in the MAR and (ii) the subMAR is unique, all valid clone trees of t are completions of the subMAR.

SubMARine is implemented in Python and can be downloaded at https://github.com/morrislab/submarine. Next to the algorithm, we provide an implementation of a depth-first search to enumerate the set of valid subMAR-completing clone trees and an upper bound on the size of this set (see Section S4.5 for a derivation of this bound).

## 4 Extended SubMARine: Clone tree reconstruction with subclonal CNAs

The extended version of SubMARine propagates inference rules like the basic version but is designed specifically to include subclonal CNAs. For example, unlike the basic version, it propagates the lost allele rule; because whether or not the lost allele constraint is satisfied depends on the choice of clone tree. Its subMAR, which we call the *extended subMAR*, defines not only the set of valid clone trees but also a set of equivalent ones and approximates the *extended maximally-constrained ancestral reconstruction problem* defined below. Two clone trees are **equivalent** if they fit the experimental data equally well and if the same set of subclonal CNAs has the same impact on the mutant copy numbers of the same set of SSMs. Given subclonal frequencies and the assignment of SSMs and clonal CNAs to subclones, as in the basic version of SubMARine, the data fit does not depend on the ancestral relationships in the clone tree [20]. However, with subclonal CNAs, ancestral relationships can influence data fit. Specifically, subclonal CNAs change the VAFs of SSMs by altering their mutant copy numbers per cancer cell but only if 1) the subclonal CNA is in a descendant subclone, 2) the SSM is in the segment affected by the CNA and 3) the SSM is on the same parental allele, i. e., it has the same *phase*, as the CNA. As such, changing the ancestral relationship between an SSM-containing subclone and a CNA-containing one, can change the fit of the clone tree to the experimentally-measured VAF data. Note that because we model the change in CNA state, rather than the absolute copy number, the data fit to the experimental-derived average copy numbers of segments is not affected by the clone tree, see Section S5.1 for details. We represent the impact of CNAs on SSMs in an impact matrix 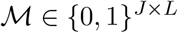, where *J* is the number of SSMs and *L* the number of CNAs:

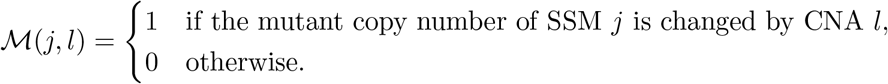

As an aside, defining 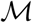 requires us to assume each SSM is unique, i.e., we make an infinite sites assumption, otherwise we would not be able to select which version of the SSM is impacted by the CNA. Given the above, if two clone trees with the same subclonal frequencies and mutation assignments imply the same impact matrix, they also have equal data fit and are thus equivalent. Note that it is possible but exceptional rare, for two clone trees to have the same data fit but not the same impact matrix (see Section S5.2 for an example).

As indicated above, a CNA changes an SSM’s mutant copy number only under specific conditions; thus the impact matrix 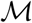 requires the presence and absence of specific ancestral relationships and SSM phases. These conditions, the **equivalence constraints**, are formally described in depth in Section S5.3 and their derived inference rules are propagated by extended SubMARine.

In the **extended clone tree reconstruction problem**, one is given a subclonal frequency matrix *ϕ*; *L* CNAs assigned to subclones, segments and parental alleles; *J* SSMs assigned to segments and subclones; as well as an impact matrix 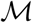; and is required to find a valid clone tree with subclonal CNA impacts that match 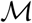. Given the input of an extended clone tree reconstruction problem *t*, the **extended maximally-constrained ancestral reconstruction problem** is to identify the pairwise ancestral relationships between subclones present in all valid clone trees that solve *t* and are thus equivalent. The **extended MAR** is the unique partial clone tree that solves this problem by defining all, and only, the ancestral relationships as well as SSM phases shared by the solution set of valid and equivalent clone trees for *t*.

Like the basic subMAR, the **extended subMAR** has three important properties for an extended clone tree reconstruction problem *t*: its defined ancestral relationships and SSM phases are a subset of those in the extended MAR, it is unique, and consequently, all valid and equivalent clone trees of t are completions of the extended subMAR (see end of Section S5.6 for more details).

As input, the extended version of SubMARine takes the subclonal frequency matrix *ϕ*, CNAs as copy number changes (i. e. gains or losses) assigned to subclones, segments and parental alleles, SSMs assigned to segments and subclones, and the impact matrix 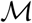 of an extended clone tree reconstruction problem (see Figure S5). Copy number changes, subclones, segments and alleles of the CNAs can be provided by subclonal CNA reconstruction methods [12, 14, 22, 25]. The impact matrix 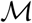 can be easily derived from an existing subclonal reconstruction – then SubMARine generalizes from one clone tree to the set of valid and equivalent ones – but in some cases it can also be inferred without a subclonal reconstruction (see Section 6). For extended SubMARine the input CNAs have to satisfy a **monotonicity restriction**, which ensures that each segment contains only copy number changes of the same direction per allele (see Section S5.4 for details). In brief, this condition guarantees that once an allele is lost, no update of undefined ancestral relationships can prevent this loss from happening (e. g. by increasing the copy number of the allele before the loss), and hence no subsequent updates to *Z* can remove conditions that used the lost allele rule to previously define an element of *Z*. This guarantees that all defined values in the subMAR set by propagating inference rules are present in the extended MAR. Note that copy neutral loss of heterozygosity (LOH) events can still be modeled because the restriction permits one of the parental alleles to be lost, and the other one to be gained.

Briefly, like the basic version of SubMARine, the extended version builds a partial clone tree by propagating the germline, generalized sum and partial tree rule and using the Subpoplar algorithm (see Figure S5). Furthermore, extended SubMARine propagates the equivalence and lost allele rules, and phases some SSMs in order to satisfy the underlying constraints (see Table S1 and Algorithm 2). In addition to user-defined ancestral relationships, the extended version of SubMARine can also take SSM phases as input. Extended SubMARine converges when no ancestral relationship or SSM phase can be propagated anymore. As SubMARine in basic version, the extended version always converges. Its result is an extended subMAR, consisting of the ancestry matrix *Z*, the possible parent matrix *τ* and the SSM phasing *π*_s_. An example of extended SubMARine and a detailed description of the algorithm, including an analysis of its polynomial runtime, can be found in Sections S5.5 and S5.6.

## 5 Results

Here, we evaluated SubMARine by applying it to simulated basic and extended clone tree reconstruction problems, thus without and with CNAs; and by applying it to data from the large, multi-sample TRACERx study [40, 41].

### 5.1 Simulated data

Section S6.1 provides a detailed description of the creation of our noise-free simulated datasets. In brief, we generated a dataset without CNAs containing 600 subclonal reconstructions, evenly divided between those with 5, 20 and 50 subclones (plus the germline); and another dataset with clonal and subclonal CNAs containing 1800 subclonal reconstructions, each with 20 subclones. The CNA-containing subclonal reconstructions are evenly divided among 9 groups of simulations where we try all combinations of the number of segments, selected from 10, 20, and 40, and the number of CNAs, selected from 10, 20, and 40. In each of the CNA-containing datasets, we randomly assigned CNAs as copy number changes to subclones, segments, and parental alleles, ensuring that a deletion is only allowed once per segment and allele. We also randomly assigned 200 SSMs to subclones, segments, and parental alleles, considering the impact of subclonal CNAs. For both types of datasets and each parameter combination, we draw 10 random subclonal reconstructions for each of 1 to 20 samples, resulting in 200 subclonal reconstructions for each parameter combination.

SubMARine constructed each subMAR (basic or extended) in less than 70 seconds using a single thread with less than one GB of memory. On average, increasing the number of samples or decreasing the number of subclones decreases uncertainty in a clone tree [35, 36]. The implied ambiguity in the subMAR solutions shows the same behavior when applied to our simulations(see Figures S6, S7, and S8). Including CNAs in our simulations further decreases uncertainty (see Figures S6 and S8) due to the additional implied ancestral constraints. Notably, in all simulations with twelve and more samples, the resulting subMAR had no undefined ancestral relationships, indicating that it had found the single clone tree solution to the reconstruction problem.

We then assessed how accurately SubMARine’s subMARs matched the actual ambiguity in the solution sets of clone trees fitting the 2400 clone tree reconstruction problems. Because each solution set is a subset of the clone trees completing the subMAR, we used a depth-first search (DFS) algorithm that incorporated the subMAR and the Subpoplar algorithm to enumerate these solution sets. Note that because not all spanning trees complete the subMAR (see Section S3.1), we do not use the Gabow-Myers-based algorithm previously employed for this task [11, 12, 19]. For 1844 of the 2400 clone tree reconstruction problems, the subMARs were completely defined, so they only had a single clone tree solution. Among the remaining 556 problems, only one of the problems predicted to have multiple solutions by SubMARine had only a single clone tree solution. So, in > 99.9% (1844/1845) of problems with a single solution SubMARine identified that solution. Of the remaining 555 problems, in 46 cases, our DFS algorithm did not complete its enumeration in less than 120h on a single thread.

For 80.4% of the 510 clone tree reconstruction problems for which we were able to fully enumerate the solution set, and that SubMARine predicts to have > 1 clone tree solution, the subMARs precisely matched the MAR. For all 2400 subMARs, we computed the recall, i.e., the proportion of the non-trivial ancestral relationships (those for subclones *k* and *k′* where 0 < *k* < *k′*) recovered from the MAR. Trivial ancestral relationships are those with which *Z* is initialized. As Figure 2 illustrates, the more constrained the clone tree reconstruction problem is, either by a higher number of samples or the presence of CNAs, the higher is the recall. With CNAs, there is 100% recall with six or more samples, without CNAs, this is true for ten or more samples.

**Fig. 2.**
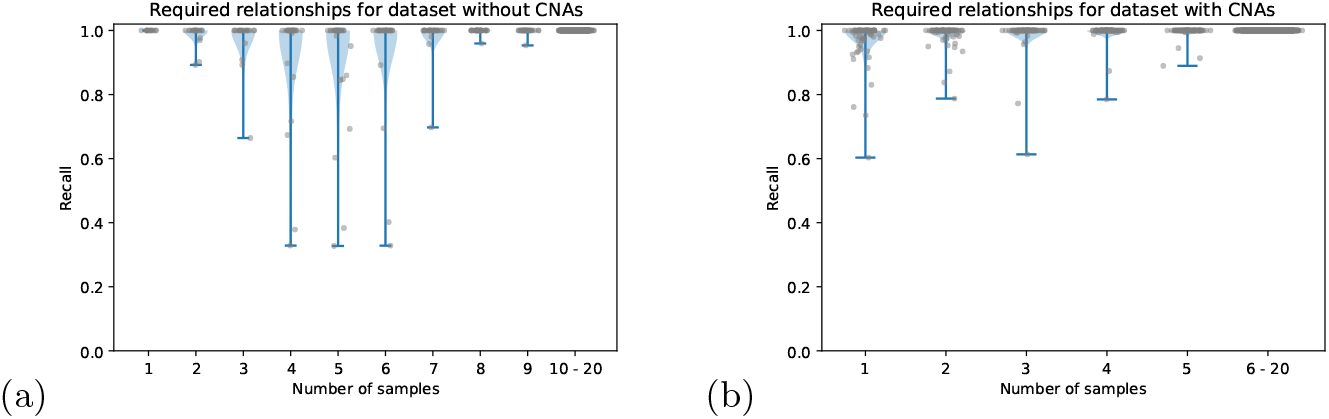
Recall of required ancestral relationships for dataset (a) without and (b) with CNAs. We computed recall based on the non-trivial ancestral relationships. Columns in (a) usually have 30 data points, columns in (b) 90. The last column in each subfigure shows all results for (a) 10 and (b) 6 and more samples since each subMAR achieved a recall of 100%. For the 46 subMARs for which the DFS could not enumerate all valid (and equivalent) completing clone trees, we did not compute the recall because we do not know the ground truth. Hence, column 1 of (a) contains only 13, column 2: 20, column 3: 21 and column 4: 26 values, and column 1 of (b) contains only 83.

As Figure 3 illustrates, it may be possible to assess when the subMAR is a perfect match to the MAR. For the dataset without CNAs, all subMARs with 5 subclones have 100% recall (Figure 3a) as do the vast majority of subMARs with less than 50 undefined relationships (Figures 3b and 3c). For the dataset with CNAs, predicting when a subMAR has 100% recall is less straightforward as there is less than perfect recall with as few as 10 undefined relationships in the subMAR (Figure 3d). However, in the CNA-containing cases, the DFS is feasible to apply for subMARs with less 50 undefined relationships as for the vast majority it was done in less than 100 seconds (see Figure S9).

**Fig. 3.**
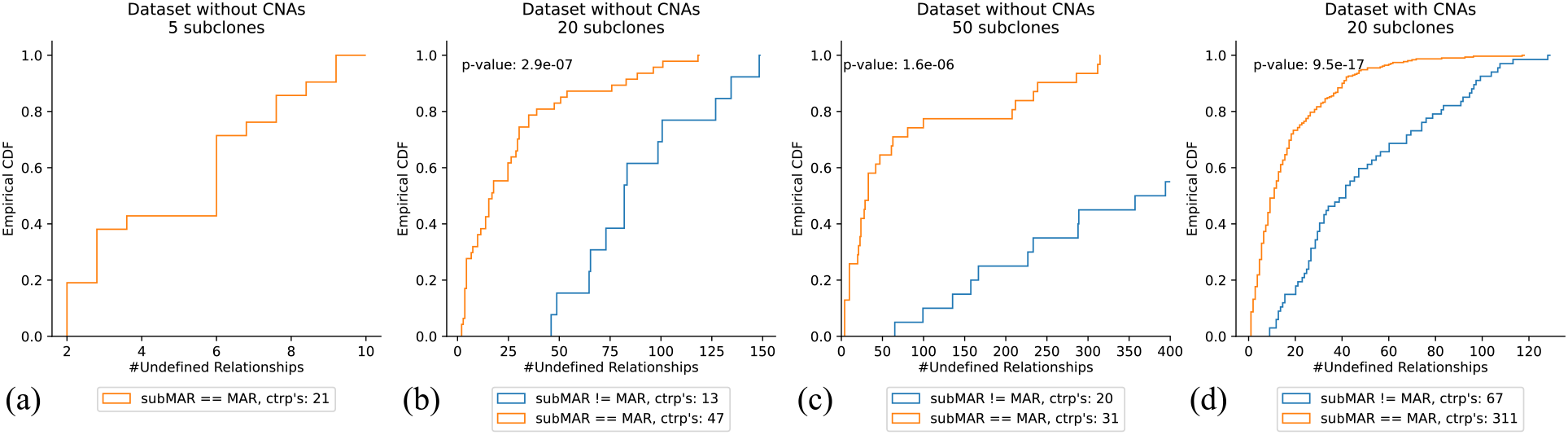
Empirical cumulative density functions (CDFs) of subMARs equaling and differing the MAR for (a)-(c) dataset without CNAs and (a) 5, (b) 20 and (c) 50 subclones, and (d) dataset with CNAs and 20 subclones. Not included are subMARs that do not contain any undefined ancestral relationships (and thus have found the single clone tree solution and equal the MAR), and those for which the DFS did not finish. The p-values are computed with a Kolmogorov-Smirnov test. (c) The empirical CDF for subMARs differing the MAR reaches the value of 1.0 at 864 undefined relationships. ctrp’s: clone tree reconstruction problems

### 5.2 TRACERx data

We next applied SubMARine to a large, multi-sample dataset drawn from the TRACERx study [41], consisting of mostly primary tumors of 100 patients with early-stage non-small-cell lung cancer (NSCLC). Previously, PyClone [28] was used for each patient to identify mutation clusters, which correspond to subclones, and CITUP [34] was used to infer clone trees by exhaustively exploring all possible trees and reporting those with the highest likelihood. In Section S6.2, we describe how we arrive at 88 patients with two to 15 subclones from two to seven tumor samples, on which we apply the basic version of SubMARine (see Table S4 in Appendix 2).

For each patient, SubMARine constructed the subMAR in less than 40 seconds on a single thread with less than one Gb of memory. For 42 patients, we did not use a noise buffer because their subclonal frequencies supported a valid partial clone tree; 37 of those have a subMAR that describes only a single tree. Figure S10 shows the five subMARs with undefined ancestral relationships. All five subMARs were identical to their MARs. In order to build a valid partial clone tree for the other 46 patients, we computed subclone- and sample-specific noise buffer sets (see Section S4.3). For 45 of these patients, the noise buffer sets could be found in polynomial time. Only for one patient (CRUK0016), an exhaustive search had to be applied; it found the MAR and the noise buffer set in less than 2 seconds. The maximum values in the noise buffer sets range from 0.01 to 0.7 (see Figure S11), with a median of 0.14. Only one patient required a buffer greater than 0.5 (see Figure S12), this could be caused by infinite sites violations [42] or an undetected CNA. With the noise buffers, SubMARine identifies 42 additional subMARs that describe a single tree. For three of the four remaining patients, SubMARine finds subMARs with one, three and seven uncertain values being perfect matches to their MARs. The MAR of the remaining patient CRUK0016 contains nine undefined values.

We next compared SubMARine’s partial clone trees with those clone trees reported in the TRACERx paper (p.31–p.174 of the Supplementary Appendix 1 of the work of Jamal-Hanjani et al. [41]). All but the trees for six patients were generated by CITUP. Full details of this comparison are provided in Table S4 in Appendix 2. CITUP exhaustively enumerates all clone trees, up to ten subclones. As such, for the three patients (CRUK0032, CRUK0062 and CRUK0065) with more than ten subclones, CITUP could not be run and the authors constructed trees manually. Note that for these three patients, SubMARine identified subMARs in less than 40 seconds. For each tree, CITUP infers a set of subclonal frequencies that are close to the input frequencies and for which the associated clone tree is valid. Trees are ranked based on how close the input and inferred frequencies are, as assessed using a likelihood function. This function is maximized when the input frequencies already support a valid clone tree. As such, for the 42 patients which did not require a noise buffer, CITUP should find the same trees as SubMARine, assuming that only the most likely trees were reported. However, for six of the 42 patients, Jamal-Hanjani et al. report more trees. None of these additional trees were valid with the unaltered frequencies (see Figure S13 for an example). In 29 of the 46 cases requiring a noise buffer, the subMAR perfectly matches the trees reported by Jamal-Hanjani et al. Of the remaining 17 cases, in twelve cases, the valid trees completing the subMAR are a subset of the reported ones, and in one case one reported and completing tree are identical but CITUP finds more trees. In the remaining four cases, there is no overlap between reported and completing trees; however, the trees differ only in up to three parent-child relationships.

## 6 Conclusion and discussion

Here we have introduced SubMARine, a polynomial-time algorithm that computes the *subMAR*, a partial clone tree that is a relatively simple, partial solution to the NP-complete problem of finding a valid clone tree for a subclonal frequency matrix *ϕ*. Despite that the subMAR is only an approximation, in almost all cases, when there is only a single clone tree solution, assuming precisely measured subclonal frequencies, SubMARine identifies it. Indeed, the subMAR only fails to capture the vast majority of the non-trivial ancestral relationships in the MAR when the reconstruction problem is severely under-constrained by the input data; and often these cases can be diagnosed by examining the subMAR. Notably, SubMARine also solves a potentially much more difficult extension of the basic clone tree reconstruction problem that includes subclonal CNAs (see also [43]). Furthermore, SubMARine permits the addition of user-defined ancestral constraints and SSM phasing, which could come from single cell or long read sequencing data. Additionally, we introduced a noise-buffered version of SubMARine to deal with inaccurate subclonal frequencies. This version finds a minimum noise buffer that is added to parental subclonal frequencies in order to prevent generalized sum rule violations and hence permits a valid partial clone tree for an input dataset.

The partial clone tree is a particularly useful summary in domains, e. g. cancer therapy, where false positive claims on the evolutionary history of a tumor can have drastic consequences. Here, a conservative assessment of uncertainty is far superior to a random or representative single clone tree solution.

Assuming precisely measured subclonal frequencies, SubMARine was able to construct the subMAR for nearly half of the TRACERx data where subclones were defined by mutation clustering. For the rest of the data, SubMARine could construct the subMAR using subclone- and sample-specific noise buffer sets. The noise-buffered version of SubMARine still requires an ordering of the subclones to initialize; the computation of this ordering does not consider the noise buffer and may be the source of differences between the solution sets reported by SubMARine and by CITUP on the TRACERx data.

Currently, SubMARine characterizes uncertainty in a clone tree assuming fixed subclonal frequencies which could lead to overconfidence in a single subMAR. Even the noise-buffered version, when working only with the minimum necessary noise buffer, basically makes this assumption. In order to account for uncertainty in subclonal frequencies, a larger noise buffer could be used. Another possibility may be to sample small amounts of noise and add these to the subclonal frequencies. Repeated multiple times, SubMARine could be applied to the different subclonal frequency sets and the resulting subMARs could be combined into a single one. One could even go one step further and add noise to the initial mutational frequencies that are input to an algorithm determining subclonal frequencies. However, because the subclones derived from different mutational frequency sets might be associated with different mutations, a mapping between subclones has to be derived. Either of these approaches may provide a principled way to identify a solution set of clone trees with nearly equivalent data fits; this would also permit use of SubMARine for datasets with low purity or low sequencing depth, for example.

An important use of SubMARine is generalizing a single clone tree – produced, e.g., through Monte Carlo sampling – to the set of equivalent clone trees. Given a clone tree, one can easily estimate a set of *ϕ* which fit the data well and satisfy the sum constraint; as well as defining the impact matrix 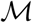. SubMARine could then identify the equivalence class of trees with equally good fits, thereby enhancing methods that give single or sampled solutions to a reconstruction problem. Indeed, assuming that a mapping between subclones could be defined between different clone trees, one could group different clone tree samples together based on their associated subMARs.

There are a number of potential further extensions of this work. It may be possible to define the impact matrix 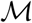 without a full subclonal reconstruction by adapting some of the pairwise comparisons technique developed in [43]. Indeed, it is possible to infer 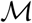 directly for subclonal CNAs that are clonal in some, but not all, samples.

A potential drawback of SubMARine is the monotonicity constraint on the subclonal CNAs; note that this constraint is both more and less limiting than the infinite allele assumption previously applied to subclonal CNAs [43]. In particular, it effectively rules out incorporating clonal whole genome duplications (WGD) that appear in many cancers. It may be possible to extend SubMARine to incorporate clonal WGDs by expanding the number of potential phases for an SSMs.

There are a number of unanswered theoretical questions raised by this work. First, it is unclear what the hardness of the MAR reconstruction problem is. Because a MAR only exists if there is at least one valid clone tree solution, it seems likely that MAR reconstruction is at least as hard as the problem of finding a single clone tree solution. However, it is not clear whether this hardness changes under the assumption that a valid clone tree exists. Neither of these two questions are addressed by SubMARine. Also SubMARine approximates the MAR but provides no guarantees about its approximate factor. It would be useful to provide such guarantees, if they exist. Or perhaps a different algorithm to generate subMARs can provide them.

SubMARine could also be viewed as an extension of methods that perform haplotyping via perfect phylogeny [44, 45]. In quadratic time, these methods solve a special case of the basic clone tree reconstruction problem, in which all elements of the subclonal frequency matrix *ϕ* are either 0, 0.5, 1. Furthermore, they provide a complete, polynomial-space summary of all valid clone trees. Their summary methods could be generalized and applied to the possible parent matrix *τ* produced by Subpoplar.

## Supporting information

**Appendix 1:** Supplement.

**Appendix 2:** Supplementary Table S4.

**Simulated data:** https://github.com/morrislab/submarine_data/archive/v1.0.zip.

## Acknowledgments

LKS was partially funded by the International DFG Research Training Group GRK 1906/1, and is now funded by a MITACS elevate postdoctoral fellowship. Part of this work was performed while LKS was affiliated with and funded by Bielefeld University and ETH Zurich. JW is funded by the NSERC CGS-D program. GR is partly funded by the “Swiss Molecular Pathology Breakthrough Platform”, funded by the ETH Special Focus Area “Personalized Health Related Technologies”, grant number #106. QM is supported by an NIH grant (P30-CA008748), an Associate Investigator Award from the Ontario Institute of Cancer Research (which partially supports LKS), a subgrant from the Canadian Centre for Computational Genomics genomics technology platform funded by Genome Canada, and holds a Canada CIFAR AI chair. We thank our reviewers, and Ben Raphael, for constructive feedback and useful suggestions.

## Author contributions

All authors devised the conceptional ideas of this work. LKS, JW and QM developed the algorithms. JW simulated the data. LKS wrote the code and performed the experiments. GR, JS and QM supervised the work. LKS and QM wrote the manuscript. All authors reviewed the manuscript.

## Supplement

### S1 Supplementary figures, tables and algorithms

**Fig. S1.**
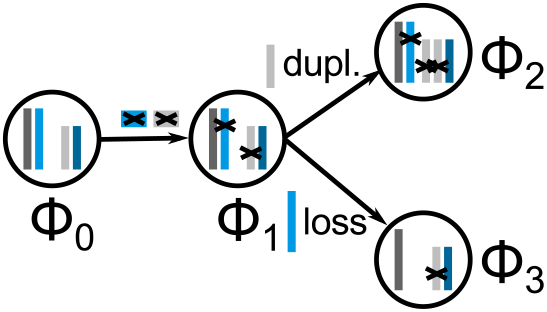
Example of a clone tree with three subclones and the germline. Subclonal frequencies are indicated with *ϕ*_0_,…,*ϕ*_3_; assuming that there are two samples given, their values could be *ϕ*_0_ = (1,1), *ϕ*_1_ = (0.9, 0.8), *ϕ*_2_ = (0.5, 0.3), and *ϕ*_3_ = (0.4, 0.35). Edges between subclones indicate ancestral relationships, with the germline being an ancestor of all subclones and subclone 1 being the ancestor of subclones 2 and 3. Colorful bars indicate alleles of different segments; here, the two alleles of two segments are shown, with segment 1 having the dark gray and the light blue alleles, and segment 2 having the light gray and dark blue alleles. Two SSMs are assigned to subclone 1, one to the blue allele of segment 1 and one to the gray allele of segment 2. The SSMs are inherited by the descendants of subclone 1. Furthermore, two CNAs are assigned to the subclones, shown as copy number changes. One copy number duplication of the gray allele of segment 2 is assigned to subclone 2, duplicating the SSM lying on it. One copy number loss of the blue allele of segment 1 is assigned to subclone 3, deleting with it the SSM of this segment.

**Fig. S2.**
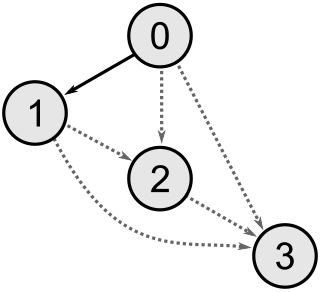
Partial clone tree where one completing clone tree is not a solution to the basic clone tree reconstruction problem *t*. Given *t* with subclonal frequency matrix *ϕ* = (1, 0.7, 0.3, 0.2)^*T*^, this partial clone tree is its MAR. Six clone trees complete the MAR, however, only five of them are valid. The clone tree in which the germline is a parent of subclones 2 and 3 does not satisfy the sum constraint and hence is not a solution to *t*.

**Fig. S3.**
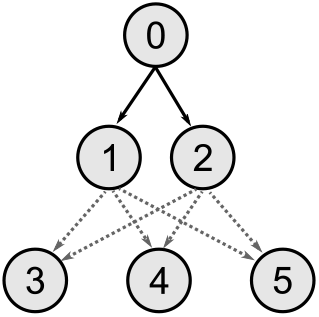
Valid partial clone tree without a valid completion. Example of a valid partial clone tree given the subclonal frequency matrix *ϕ* = ((1.0,1.0), (0.6, 0.6), (0.4, 0.4), (0.39, 0.37), (0.38, 0.38), (0.37, 0.39))^T^. Subclones 1 and 2 are definite children of the germline. Subclones 1 and 2 do not have definite children because their ancestral relationships to subclones 3, 4 and 5 are undefined. In a completion without undefined relationships, either subclone 1 or 2 would have to have two definite children. However, given the frequencies in *ϕ*, subclones 1 and 2 can have only one definite child without violating the generalized sum constraint. Thus, there exists no valid full completion of this valid partial clone tree.

**Fig. S4.**
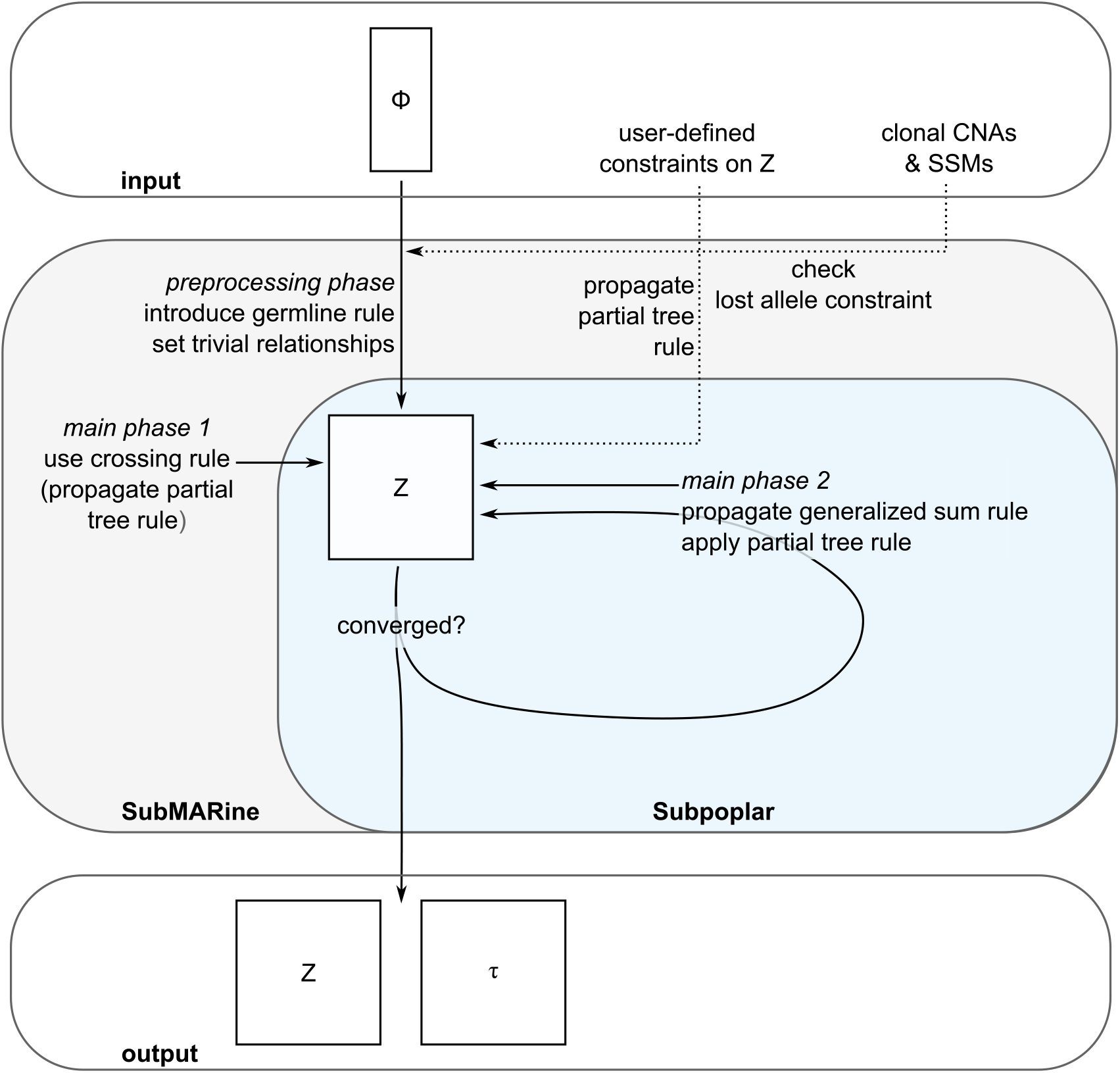
Overview of SubMARine in basic mode. The basic version of SubMARine takes the subclonal frequency matrix *ϕ* as input to build the ancestry matrix *Z*. In a preprocessing phase, the germline rule is introduced by setting *Z*(0, *k*) = 1 for all *k* > 0. Also, all trivial relationships are set to 0 (*Z*(*k, k′*) = 0 for *k′ ≤ k*) as a consequence of the generalized sum constraint. Then, the main phase starts by using the crossing rule (Equation 9), which also follows from the generalized sum constraint. The generalized sum rule itself and the partial tree rule are propagated by using Subpoplar until the ancestry matrix converged and no more relationships can be defined. Then, SubMARine outputs the ancestry matrix *Z* together with the possible parent matrix *τ*, created by Subpoplar. When the user defines additional constraints on *Z*, these are also input to SubMARine. They are applied after the preprocessing phase, followed by a propagation of the partial tree rule. This rule is also propagated now when using the crossing rule. The reason is that with the entries set by the user, *Z* can contain 1’s in other positions than the first row, possibly requiring updates of undefined relationships. Without user-defined constraints on *Z*, 1’s in other rows can be set only in Subpoplar, hence the partial tree rule needs to be applied only at that stage. When the user provides clonal CNAs and SSMs as input, the lost allele constraint is checked before starting the preprocessing phase. Whenever a constraint cannot be satisfied, SubMARine terminates and indicates which subclonal relationship led to the conflict.

**Fig. S5.**
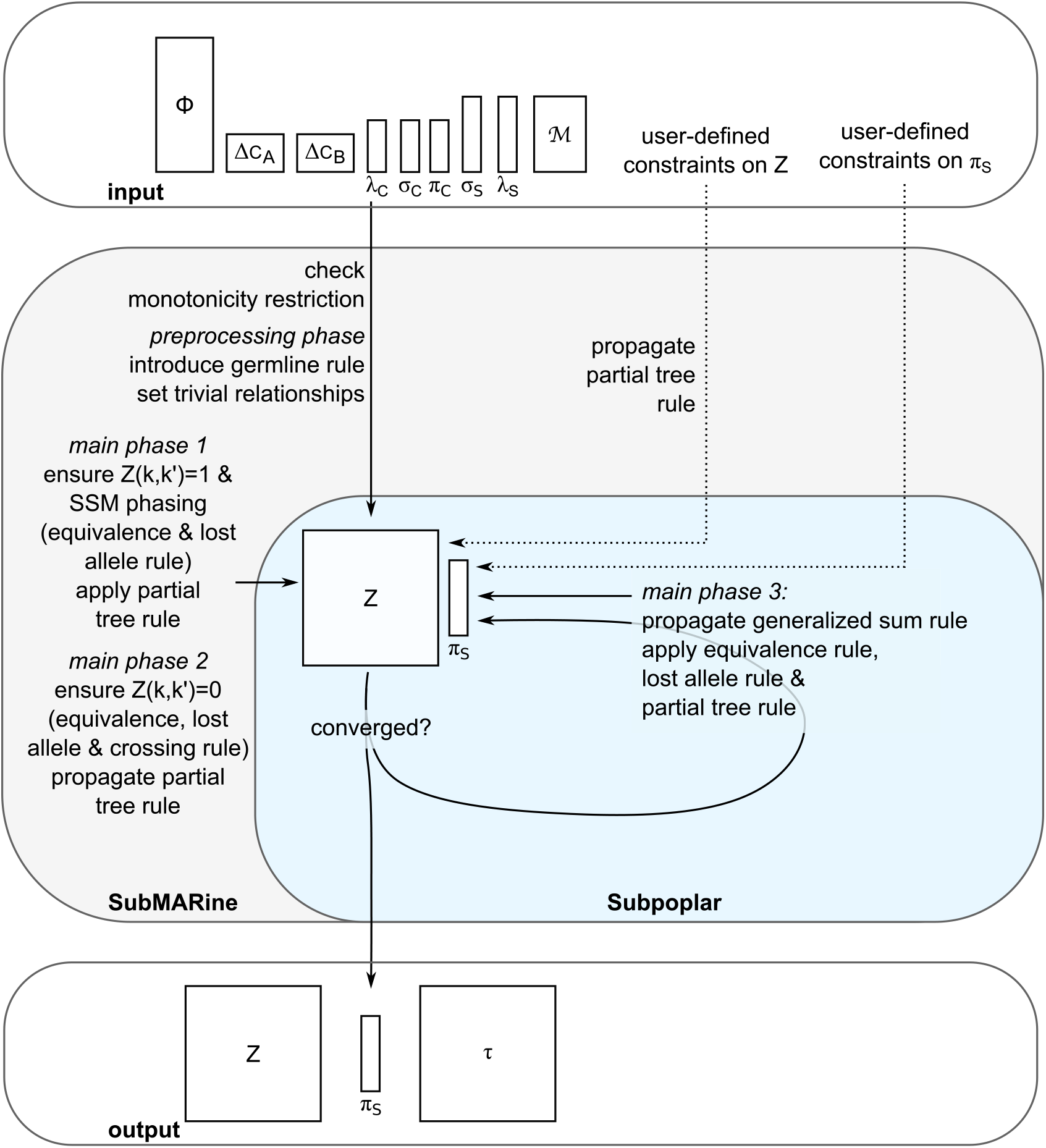
Overview of SubMARine in extended mode. The extended version of SubMARine takes the subclonal frequency matrix *ϕ*, CNAs as copy number changes in the matrices *ΔC_A_* and *ΔC_B_*, assigned to subclones, segments and parental alleles in the vectors λ_c_, *σ*_c_ and *π*_c_, SSMs assigned to segments and subclones in the vectors *σ*_s_ and λ_s_, and the impact matrix 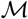 as input to build the ancestry matrix *Z* and the SSM phasing vector *π*_s_. At first, the monotonicity restriction is checked to hold on the CNAs. Then, in the preprocessing phase, the germline rule is introduced and trivial relationships (*Z*(*k, k′*) = 0 for *k′ ≤ k*) are set. Afterwards, SubMARine starts the main phase, ensuring that the partial tree rule is applied each time a relationship is updated. First, the equivalence rule based on Equation 13 is propagated, leading to 1’s in *Z*, together with those equivalence and lost allele rules that update SSM phasing. Second those equivalence and lost allele rules that lead to 0’s in *Z* and the crossing rule are used. Third, the general sum rule is propagated with Subpoplar, which also applies the equivalence, lost allele and partial tree rules whenever necessary. The method converges, when no more subclonal relationships and SSM phases can be updated. The output consists of the ancestry matrix *Z*, the SSM phasing vector *π*_s_ and the possible parent matrix *τ*, created by Subpoplar. The user can also define additional constraints on *Z* and on *π*_s_. Both types of constraints are applied after the preprocessing step and before the main phase starts. When user-constraints on *Z* are set, the partial tree rule is already propagated before the main phase. Whenever a constraint cannot be satisfied, SubMARine terminates and indicates what led to the conflict.

**Table S1.** Overview of inference rules derived from the lost allele and equivalence constraints. For explanation of relative copy numbers *ΔC_A_* and *ΔC_B_*, CNA subclone, segment and phase assignments, λ_c_, *σ*_c_, and *π*_c_, and SSM subclone, segment and phase assignments, λ_s_, *σ*_s_, and *π*_s_, see Section S2; and for impact matrix 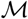 see Section 4. *Z* is the ancestry matrix. Indices *k* and *k′* refer to subclones ordered by their average frequencies, *j* to SSMs and *l* to CNAs. The function *ρ*(*α*) = *β* takes an allele *α* as input and returns the opposite allele *β*.

| inference rule | functional description | impact | application |
| --- | --- | --- | --- |
| lost allele rules |  |  |  |
| i) | $Z_{t+1} \leftarrow f_{lost_{z0}}(\Delta C_A, \Delta C_B, \lambda_s, \lambda_c, \sigma_s, \sigma_c, \pi_c, \pi_{s_t}, Z_t)$ | $Z(k, k') = 0$ (see Equations 4, 6, and 7) | once, and then every time a new 1 is set in $Z$ and an SSM got phased |
| ii) | $\pi_{s_0} \leftarrow f_{lost_{pha}}(\Delta C_A, \Delta C_B, \lambda_s, \lambda_c, \sigma_s, \sigma_c, \pi_c, Z_t)$ | $\pi_s(j) = \rho(\pi_c(l))$ (see Equation 8) | once, and then every time a new 1 is set in $Z$ |
| equivalence rules |  |  |  |
| i) | $Z_0 \leftarrow f_{eq_{z1}}(\mathcal{M}, \lambda_s, \lambda_c)$ | $Z(k, k') = 1$ (see Equation 13) | once |
| ii) | $\pi_{s_0} \leftarrow f_{eq_{samepha}}(\mathcal{M}, \lambda_s, \lambda_c, \pi_c)$ | $\pi_s(j) = \pi_c(l)$ (see Equation 14) | once |
| iii) | $\pi_{s_0} \leftarrow f_{eq_{difpha}}(\mathcal{M}, \lambda_s, \lambda_c, \pi_c, Z_t)$ | $\pi_s(j) = \rho(\pi_c(l))$ (see Equation 15) | once, and then every time a new 1 is set in $Z$ |
| iv) | $Z_{t+1} \leftarrow f_{eq_{z0}}(\mathcal{M}, \Delta C_A, \Delta C_B, \lambda_s, \lambda_c, \sigma_s, \sigma_c, \pi_c, \pi_{s_t}, Z_t)$ | $Z(k, k') = 0$ (see Equations 16, 17, and 18) | once, and then every time a new 1 is set in $Z$ and an SSM got phased |

#### Algorithm 2 Functional description of the SubMARine algorithm in extended mode

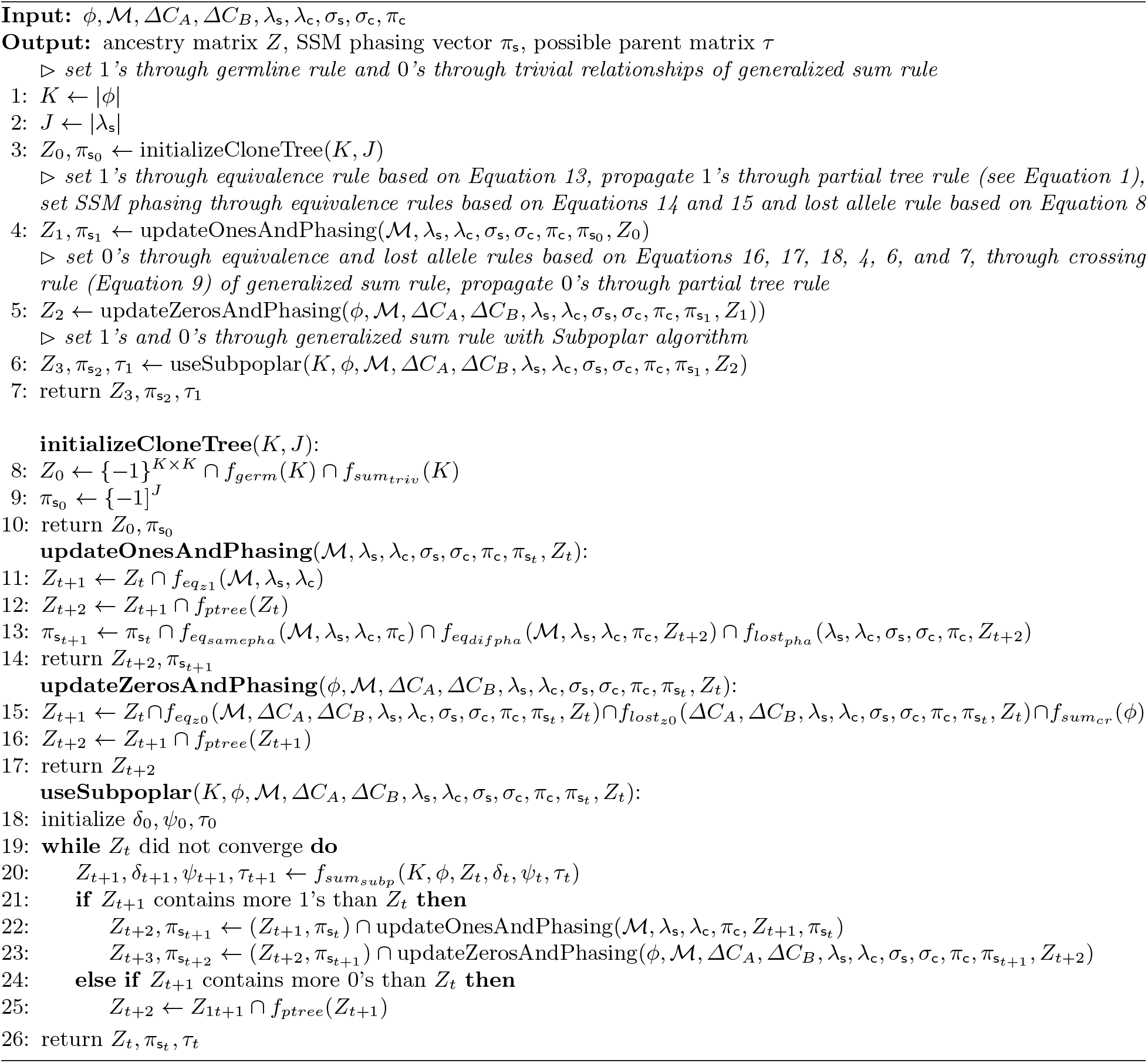

**Fig. S6.**
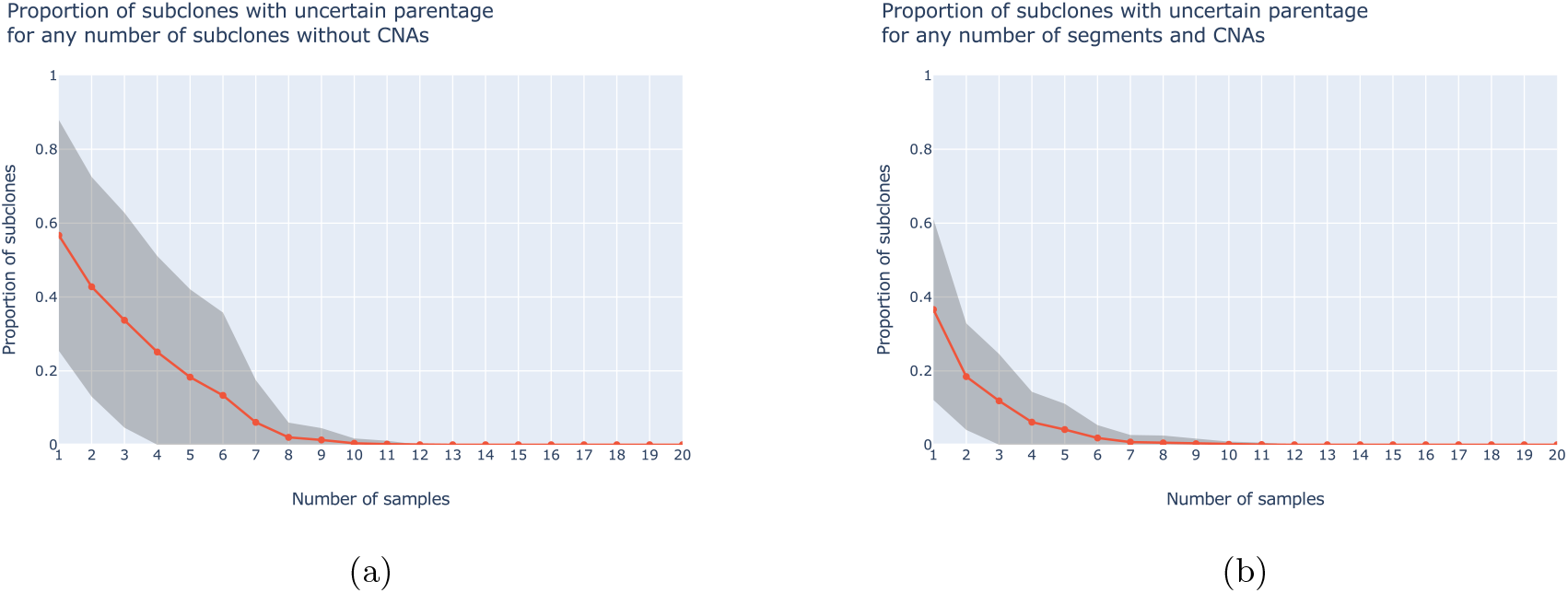
Proportion of subclones with uncertain parentage for (a) dataset without CNAs and (b) dataset with CNAs. A subclone has uncertain parentage when it has multiple possible parents in the possible parent matrix *τ*. Line show mean and gray area standard deviation.

**Fig. S7.**
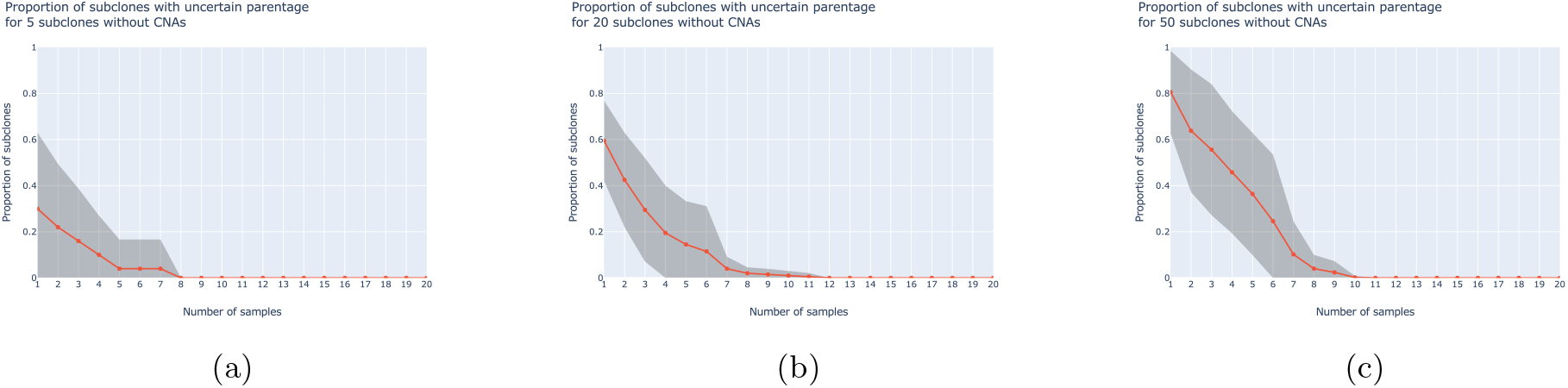
Proportion of subclones with uncertain parentage for dataset without CNAs containing (a) 5, (b) 20 and (c) 50 subclones. A subclone has uncertain parentage when it has multiple possible parents in the possible parent matrix *τ*. Line show mean and gray area standard deviation.

**Fig. S8.**
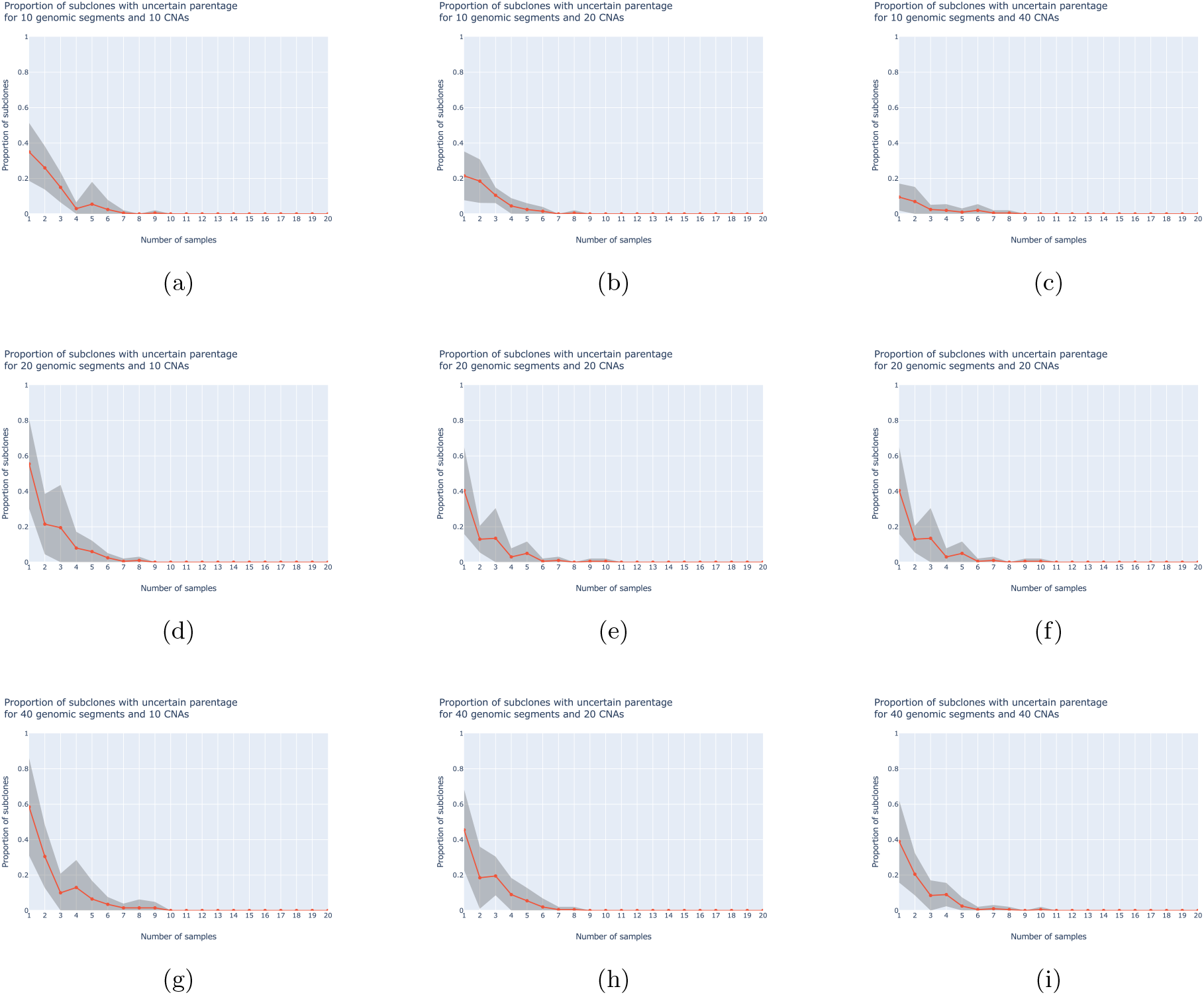
Proportion of subclones with uncertain parentage for dataset with CNAs containing 20 subclones and different numbers of CNA events and segments. (a)-(c) 10 segments, (d)-(f) 20 segments, (g)-(i) 40 segments, (a), (d), (g) 10 CNAs, (b), (e), (h) 20 CNAs, (c), (f), (i) 40 CNAs. A subclone has uncertain parentage when it has multiple possible parents in the possible parent matrix *τ*. Line show mean and gray area standard deviation.

**Fig. S9.**
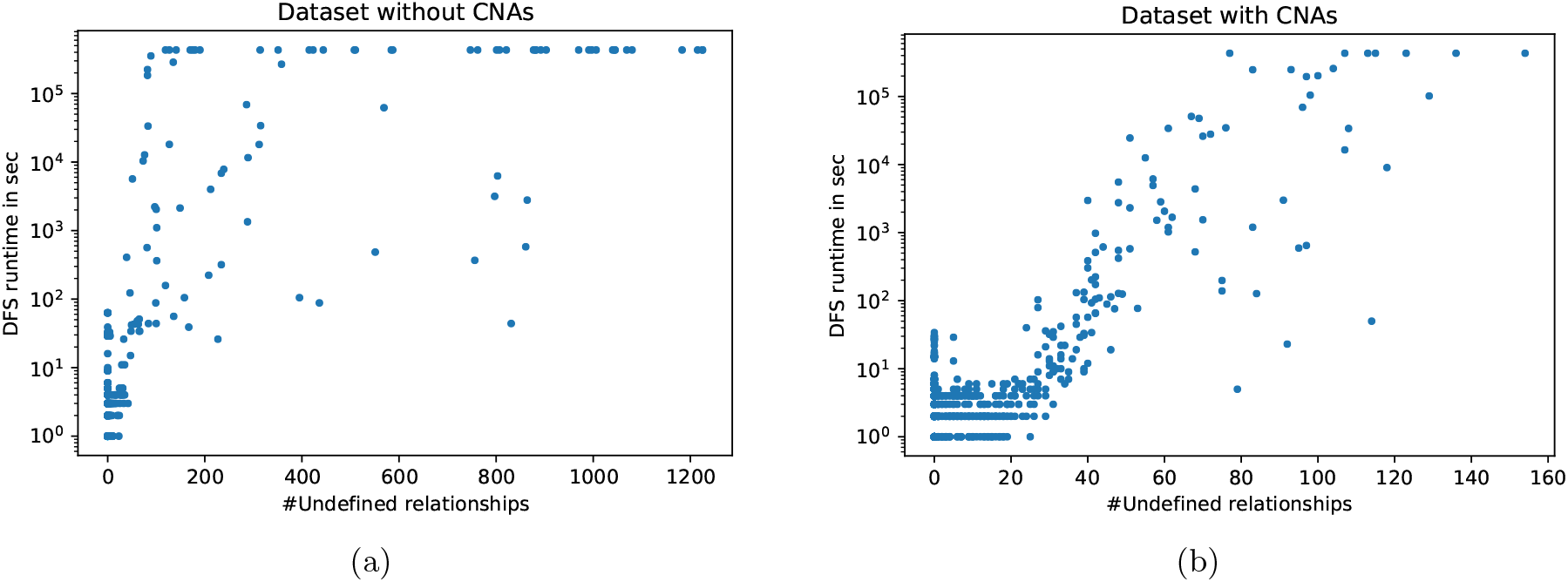
Runtimes of the depth-first search (DFS) to enumerate all valid (and equivalent) clone trees completing a subMAR, sorted by the number of undefined ancestral relationships in the subMARs. We terminated searches exceeding a maximal runtime of 120 h. We used two versions of the DFS to enumerate clone trees for different subMARs. The first version is a naïve, recursive one and the second version is an improved, iterative and also faster one, which we provide with SubMARine. Hence, if using the second version to enumerate the clone trees of all subMARs, the overall runtime could be improved. Note that for all subMARs on which the search did not termindate in 120 h, we already used the faster version.

**Fig. S10.**
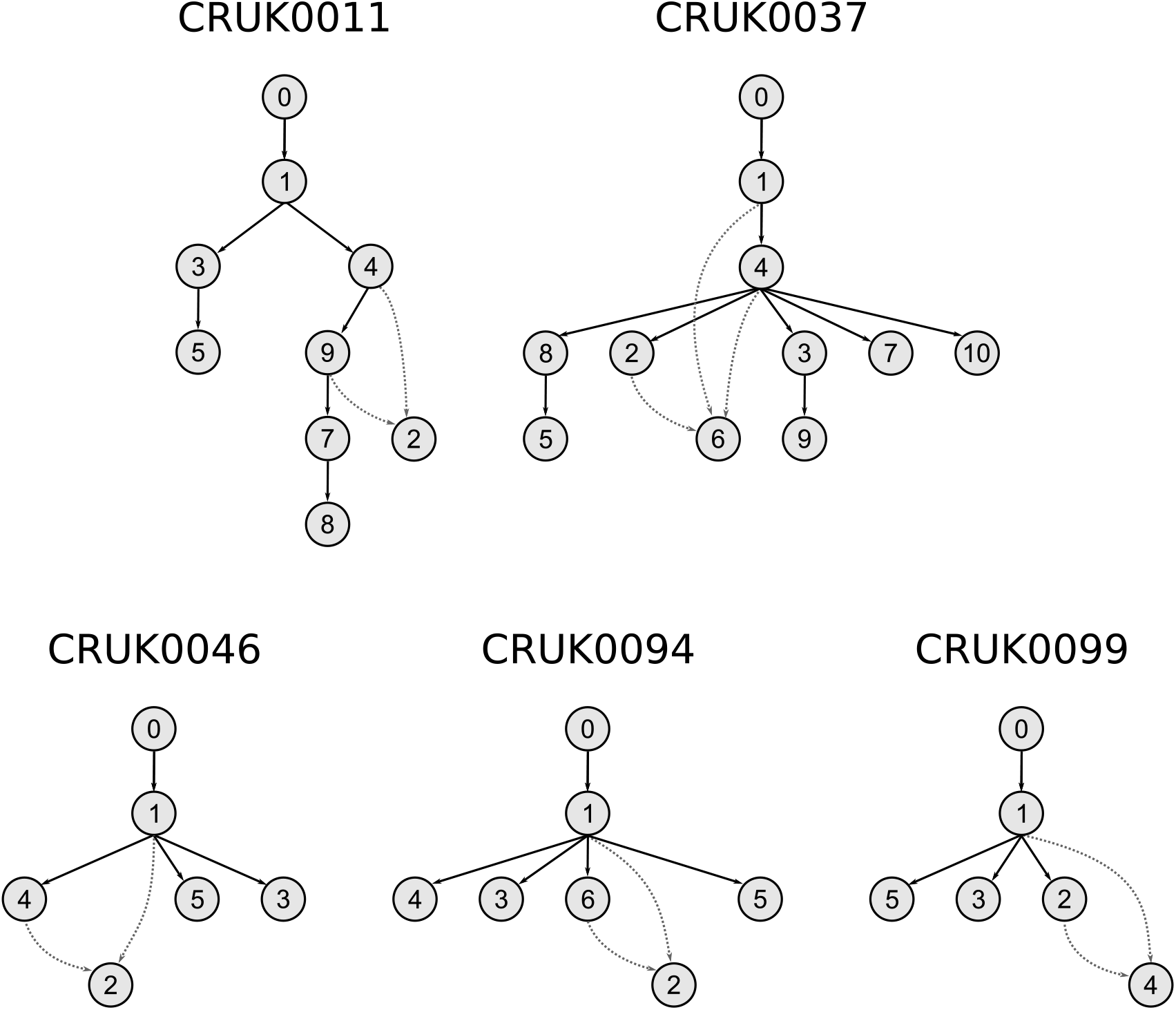
subMARs for five patients from the TRACERx cohort. Shown are the subMARs that contain undefined ancestral relationships. They are identical to their MAR. Subclonal indices are taken from the TRACERx mutation clusters.

**Fig. S11.**
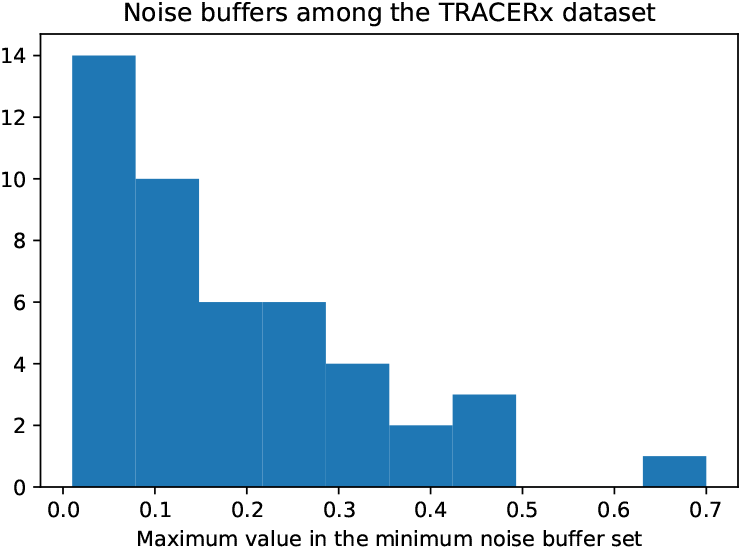
Maximum values in the minimum noise buffer sets for 46 patients of the TRACERx cohort.

**Fig. S12.**
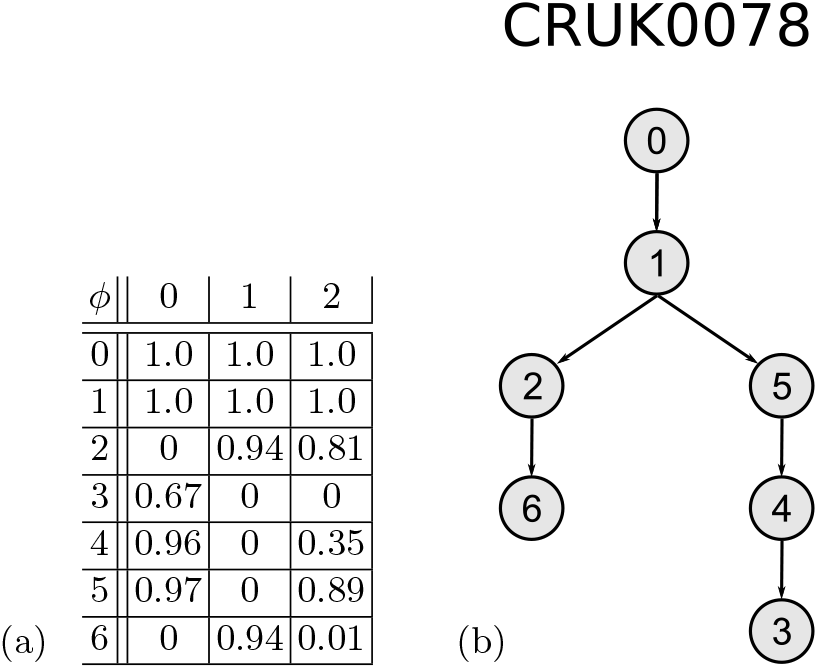
(a) Subclonal frequencies and (b) partial clone tree built by SubMARine for patient CRUK0078 of the TRACERx study. Subclonal indices are taken from the TRACERx mutation clusters. Both subclones 2 and 5 are children of subclone 1. However, they have a subclonal frequency of 0.81 and 0.89, respectively, in sample 2. Hence, a noise buffer of 0.7 is necessary.

**Fig. S13.**
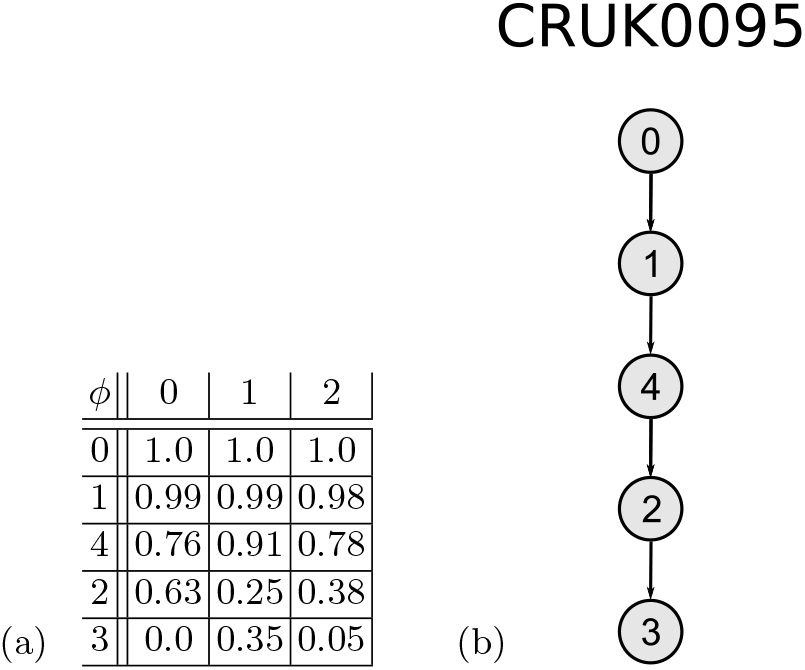
(a) Subclonal frequencies and (b) one clone tree built by CITUP in the TRACERx study for patient CRUK0095. Subclonal indices are taken from the TRACERx mutation clusters. Given the shown subclonal frequencies and the clone tree, the sum constraint is not satisfied because *Z*(2, 3) = 1 although *ϕ*(2,1) < *ϕ*(3,1). Hence, CITUP must have inferred other subclonal frequencies.

### S2 Details on the lost allele constraint

The lost allele constraint ensures that no mutation, SSM as well as CNA, gets assigned to an allele already deleted completely. In order to formulate this constraint, we need to define for each of the *L* CNAs the segment on which it occurs, and the subclone and parental allele it is assigned to. For these features, we use the vectors *σ*_c_ ∈ {0, 1,…, *I* − 1}^*L*^, λ_c_ ∈ {1, 2,…, *K* − 1}^L^ and *π*_c_ ∈ {*A, B*}^*L*^, respectively, where *I* is the number of segments, *K* is the number of subclones including the germline, and *A* and *B* are the two parental alleles. We call the alleles simply *A* and *B* because often it is not possible to determine which alleles are maternal or paternal. Note that alleles across segment boundaries are not necessarily the same; thus the A alleles of two segments do not have to be inherited both from either mother or father but one can come from mother and one from father. This is because mutations are phased only locally within one segment and not globally across all segments.

We represent CNAs as *relative* copy numbers, thus as copy number changes, and not as absolute copy numbers. Advantages of this representation are described in [22]. We store the direction and magnitude of the copy number changes for each allele in each segment *i* and subclone *k* in the matrices *ΔC_A_* and 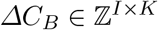 as follows:

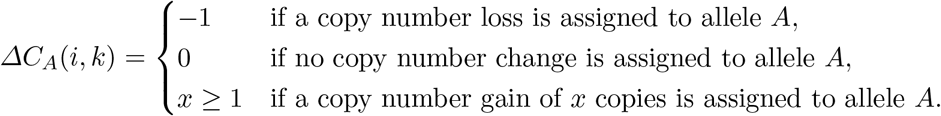

The matrix *ΔC_B_* is defined analogously for allele *B*. The normal copy number of an allele that is not influenced by copy number changes is 1.

For all *J* SSMs, the segment, subclonal and parental allele assignments are stored in the vectors *σ*_s_ ∈ {0, 1,…, *I* − 1}^*J*^, λ_s_ ∈ {−1, 1, 2,…, *K* − 1}^*J*^ and *π*_s_ ∈ {*A, B*, − 1}^*J*^, respectively. A negative value in the vectors λ_s_ and *π*_s_ indicates that the entry is undefined and the SSM is not assigned to a subclone or allele. We call an SSM *unphased* if it is not assigned to an allele.

Given the mutation assignment information, we can now formally formulate the lost allele constraint with five equations as follows:

1. If both subclones *k* and *k′* lose the same allele in the same segment and have no copy of this allele left, they cannot be in an ancestral-descendant relationship:

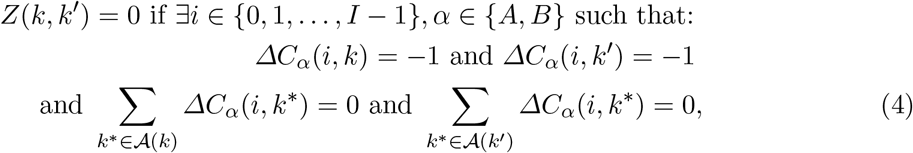

where *Z* is the ancestry matrix defined in Section 3 and where the function 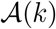 returns all ancestors of subclone *k*:

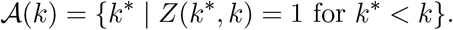
2. If all copies of an allele are lost in a segment of subclone *k*, its copy number cannot be changed in descendant subclones. Thus, subclone *k* cannot be the ancestor of subclone *k′* that contains a copy number change of this allele in the same segment:

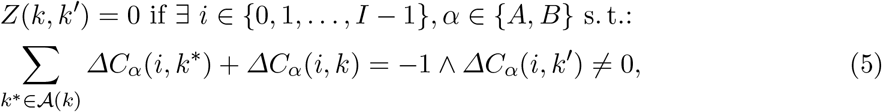
3. If subclone *k* lost all copies of one allele, it cannot be the ancestor of subclone *k′* that has at least one SSM that is phased to this allele in the same segment:

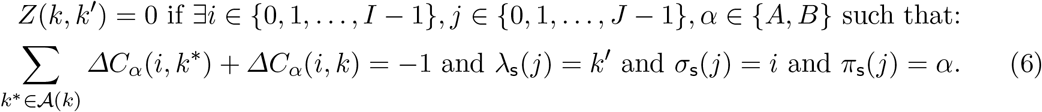
4. If subclone *k* lost all copies of both alleles in one segment, it cannot be the ancestor of subclone *k′* that has at least one SSM in the same segment:

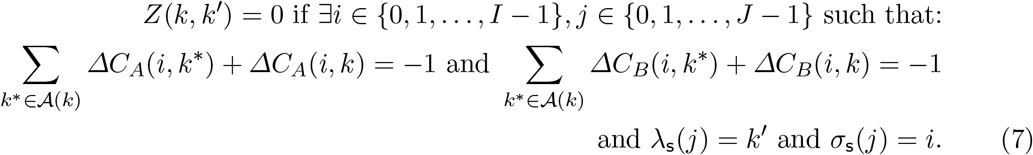
5. If subclone λ_s_(*j*) or an ancestral subclone loses all copies of an allele in segment *σ*_s_(*j*), the SSM *j* needs to be phased to the opposite allele:

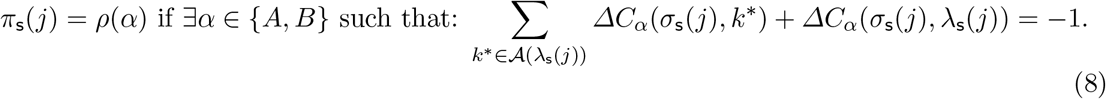

### S3 Details on partial clone trees

#### S3.1 Ancestry graphs and partial clone trees

Ancestry graphs are DAGs where the vertices represent subclones and an edge goes from subclone *k* to subclone *k′* if *ϕ*(*k, n*) ≥ *ϕ*(*k′, n*) for all samples *n* ∈ {0, 1,…, *N* − 1} and if *k′* does not contain any mutation that is already lost in *k* [11, 12, 19]. Every ancestry graph can be represented as a partial clone tree where *Z*(*k, k′*) = −1 if an edge connects *k* to *k′*, and where *Z*(*k, k′*) = 0 otherwise. To convert a partial clone tree into an ancestry graph, an edge is drawn from subclone *k* to *k′* if *k* is a possible parent of *k′* (see Definition 1). However, not every partial clone tree can be represented as an ancestry graph. It is not possible when a subclone *k* has a possible parent that has a possible parent *k\** that is not an ancestor of *k* (*Z*(*k*,k*) = 0, see Figures S14a and S14b). Ancestry graph methods enumerate clone trees as spanning trees. This approach is not intuitive with partial clone trees because not every spanning tree completes the ancestry matrix *Z* (see Figures S14a and S14c).

#### S3.2 Details on the partial tree constraint

A tree has the following two properties. First, its ancestral relationships are transitive, meaning that if node *k* is an ancestor of node *k′* and node *k′* is an ancestor of node *k″*, then node *k* also has to be an ancestor of node *k″*. Second, each node except the root, has exactly one parent. Thus, if nodes *k* and *k′* are both ancestors of node *k″*, then either node *k* has to be an ancestor of node *k′* or vice versa. Because both properties involve triplets of nodes, which correspond to our subclones, an ancestry matrix describes a tree if both properties are true for all triplets of entries. Below, we combine these two properties in the partial tree constraint.

An entry of the ancestry matrix *Z* can take three different values, leading to 27 different triplet combinations. Assuming that the subclones are sorted in decreasing order of their average sub-clonal frequencies, two combinations without undefined values violate the two tree properties and six combinations with undefined values have the potential to violate one of the two tree properties depending on whether the undefined value is set to 1 or 0 (see Table S2). Observing from enumeration of all possible combinations, all violations have in common that *Z*(*k′, k″*) = 1 and that *Z*(*k, k′*) and *Z*(*k, k″*) have different values. Thus, we can conclude the partial tree constraint:

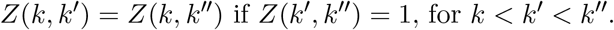

**Table S2.** Special ancestral relationship combinations of three subclones. Relationships for subclones fc, *k′* and *k″*, with 0 ≤ *k* < *k′* < *k″* < *K*, are shown in an excerpt of the ancestral matrix *Z* and with a graphical representation. A solid black edge indicates an ancestral-descendant relationship, a gray dashed edge indicates an undefined relationship and no edge between two nodes indicates no ancestral-descendant relationship. The (potentially) violated tree property is indicated for each combination.

|  | ancestral relationships | graphical representation |  |
| --- | --- | --- | --- |
| 1. | $\begin{array}{cc} & k' & k'' \\ k & 0 & 1 \\ k' & & 1 \end{array}$ | | violates single parent property |
| 2. | $\begin{array}{cc} & k' & k'' \\ k & 1 & 0 \\ k' & & 1 \end{array}$ | | violates transitivity property |
| 3. | $\begin{array}{cc} & k' & k'' \\ k & -1 & 1 \\ k' & & 1 \end{array}$ | | would violate single parent property if $Z(k, k') = 0$ |
| 4. | $\begin{array}{cc} & k' & k'' \\ k & 1 & -1 \\ k' & & 1 \end{array}$ | | would violate transitivity property if $Z(k, k'') = 0$ |
| 5. | $\begin{array}{cc} & k' & k'' \\ k & 1 & 0 \\ k' & & -1 \end{array}$ | | would violate transitivity property if $Z(k', k'') = 1$ |
| 6. | $\begin{array}{cc} & k' & k'' \\ k & 0 & 1 \\ k' & & -1 \end{array}$ | | would violate single parent property if $Z(k', k'') = 1$ |
| 7. | $\begin{array}{cc} & k' & k'' \\ k & 0 & -1 \\ k' & & 1 \end{array}$ | | would violate single parent property if $Z(k, k'') = 1$ |
| 8. | $\begin{array}{cc} & k' & k'' \\ k & -1 & 0 \\ k' & & 1 \end{array}$ | | would violate transitivity property if $Z(k, k'') = 1$ |

**Fig. S14.**
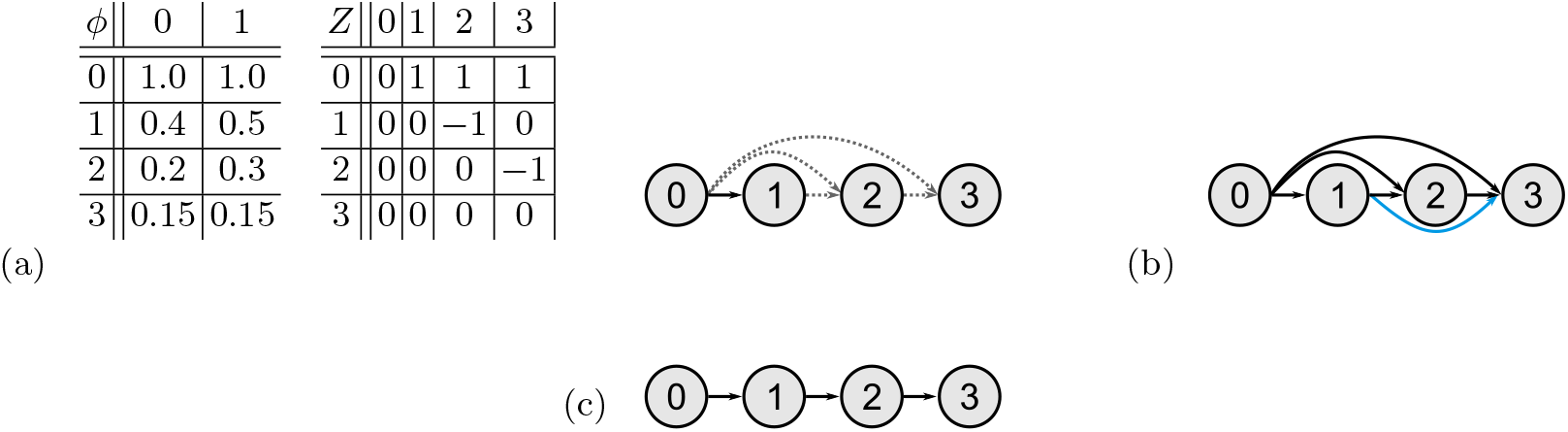
Example of (a) a partial clone tree, (b) an ancestry graph and (c) a possible spanning tree. (a) Subclonal frequency matrix *ϕ*, ancestry matrix *Z* and corresponding clone tree for an example with three subclones. Subclones 2 and 3 have two possible parents but subclone 1 is not an ancestor of subclone 3 (*Z*(1, 3) =0). We assume that no mutations get lost in this example. (b) Only the black edges would be drawn in the ancestry graph when converting the partial clone tree of (a). According to the definition of an ancestry graph however, the blue edge also needs to be present. Hence, the given partial clone tree cannot be transformed into a proper ancestry graph. (c) Spanning tree found in the partial clone tree. However, this spanning tree does not complete the ancestry matrix *Z* because *Z*(1, 3) = 0 and here subclone 1 is an ancestor of 3.

#### S3.3 Lost allele constraint for partial clone trees

The five Equations 4–8, which formulate the lost allele constraint for a complete clone tree, depend on the mutation assignments to segments, parental alleles and subclones, as well as on the definite ancestors of some subclones. Whether there is a *possible* ancestor *k*° of a subclone *k* with *Z*(*k°, k*) = −1 in a partial clone tree does not influence the lost allele constraint because as long as subclone *k*° is not a definite ancestor of subclone *k*, its copy number changes have no influence on the allele specific copy numbers of subclone *k*. Hence, the lost allele constraint does not have to be adapted to be used for a partial clone tree.

#### S3.4 Valid MAR per construction

Given all valid clone trees of a basic clone tree reconstruction problem t, constructing its MAR is trivial. All ancestral relationships that are the same across all clone trees are kept and the ones that differ are set to undefined values. The resulting partial clone tree is always valid. If it was invalid because defined ancestral relationships would violate validity constraints, then this violation would already appear in all clone trees the MAR was constructed from, thus these could not have been valid in the first place. If it was invalid because undefined values would violate the constraints, then in order to satisfy these constraints only one of the defined values would be possible. Hence, all valid clone trees would have to contain this value and consequently, it would not be undefined in the MAR. Therefore, the MAR is valid per construction.

### S4 Details on SubMARine

#### S4.1 SubMARine in basic mode

We now describe in more detail how SubMARine approximates the maximally-constrained ancestral reconstruction problem in basic mode and analyze its runtime. *K* is the number of subclones including the germline and N is the number of samples.

##### Algorithm 3 Pseudocode of the SubMARine Algorithm in Basic Mode

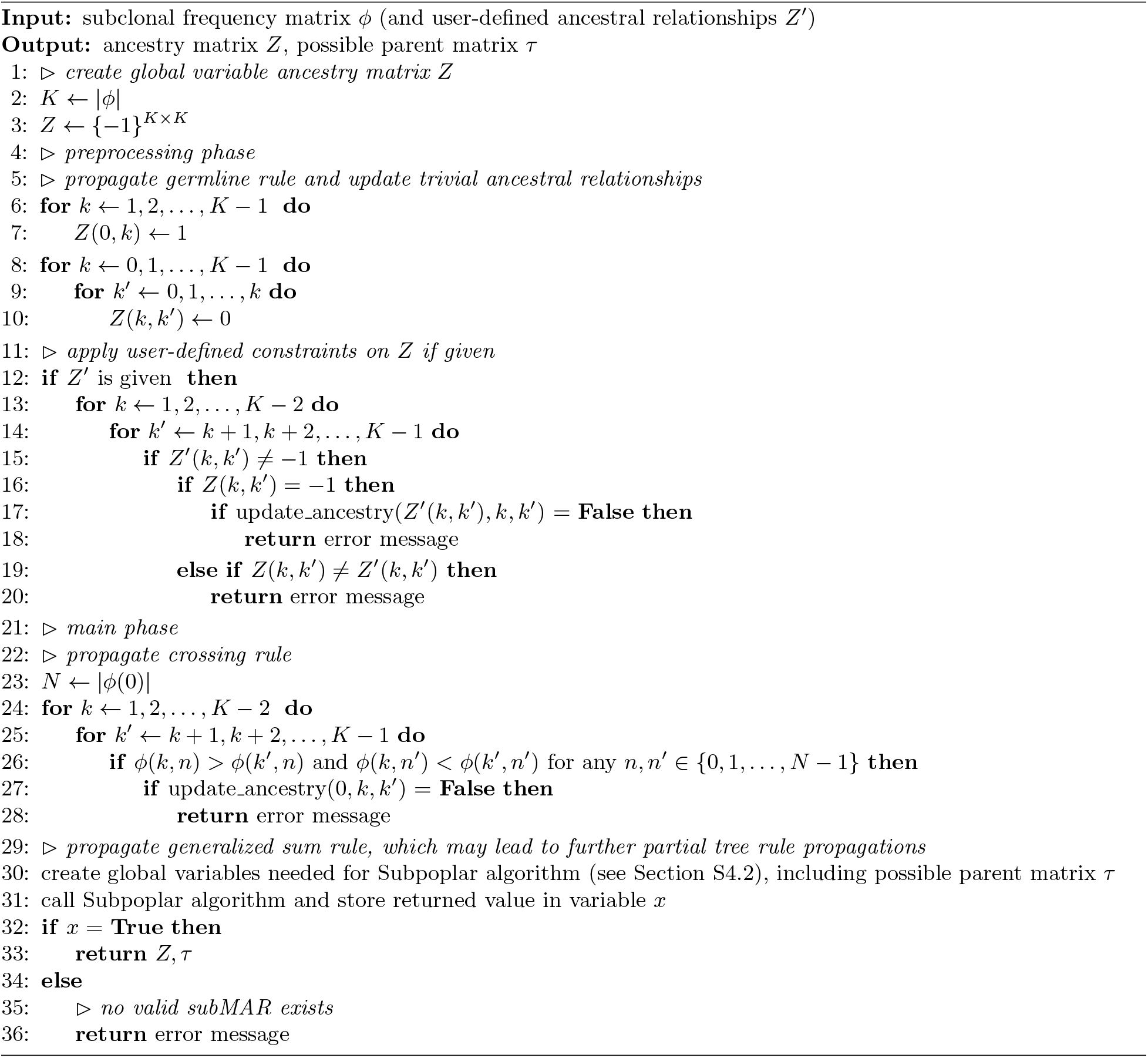

In basic mode, SubMARine (see Algorithm 3) takes only the subclonal frequency matrix *ϕ* for *K* − 1 subclones and the germline as input (see Figure S4). These subclones are sorted in decreasing order of their average subclonal frequencies across samples. If two subclones *k* and *k′* have the exact same subclonal frequencies across all samples, we arbitrarily chose one to have a lower index than the other, hence we pose a partial order also on these subclones. At first, SubMARine creates an ancestry matrix *Z* where all relationships are initially undefined 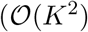 time, line 3 of Algorithm 3). It then begins with a small preprocessing phase, propagating the germline rule (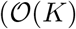 time, lines 6 and 7), and updating trivial relationships, which are *Z*(*k,k′*) = 0 with *k′ ≤ k* as a consequence of the generalized sum constraint and the ordering of the subclones 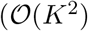 time, lines 8-10). If user-defined ancestral relationships are given, they are applied, followed by a propagation of the tree rules (lines 12 to 20 in Algorithm 3, and Algorithm 4). If we do not consider the relationship updates, this is done in 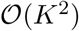 time. Considering propagating the partial tree rules lead to 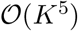 time. This is because each updated relationship can influence up to *K* other relationships, which have to be checked. Each of the influences relationships, can lead to further relationship updates. However, since each of the *K*^2^ relationship is updated at most once, the number of total updates and hence relationship propagations is limited.

##### Algorithm 4 update_ancestry(*υ, k, k′*)

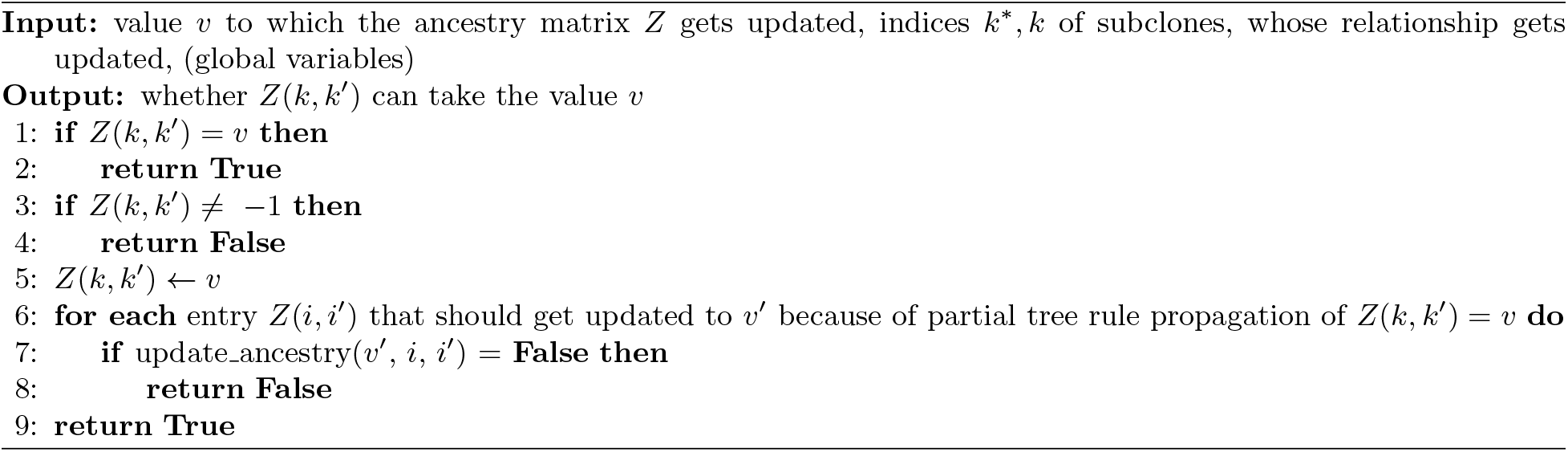

Next, the main phase starts, propagating first the crossing rule [15] as another consequence of the generalized sum constraint. This rule states that two subclones *k* and *k′* cannot be on the same branch of the clone tree if *k* has a higher subclonal frequency than *k′* in sample *n* but a lower one in sample *n′*. Because of the order of subclones and the trivial relationships, we know that *Z*(*k′,k*) = 0, and hence can implement the crossing rule as:

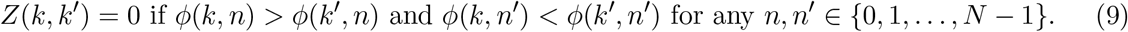

Propagating the crossing rule (Equation 9) naïvley (lines 24-28) without considering relationship updates takes 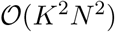 time. However, because of the ordering of the subclones, we know that the frequency of subclone *k*, with *k < k′*, is higher than or equal to the frequency of subclone *k′* in at least one sample. Thus, by checking only whether subclone *k* has a lower frequency than subclone *k′* in at least one sample, we can reduce the runtime of the crossing rule to 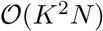 time. Its runtime when considering the propagation of the partial tree constraints is 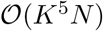. Afterwards, the generalized sum rule derived from the generalized sum constraint (Equation 3) is propagated by applying Subpoplar, which also propagates the partial tree rule. In Section S4.2, we present this algorithm, which also creates and updates a possible parent matrix *τ*, indicating the possible parents for each subclone. Furthermore, we show that Subpoplar has a runtime of 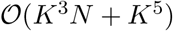, which becomes the overall runtime of SubMARine.

At the end, SubMARine returns the subMAR, consisting of the ancestry matrix *Z* and the subclonal frequency matrix *ϕ*, as well as the possible parent matrix *τ*.

It is possible for a user to define relationships for subclones. These relationships are set after the initial ancestry matrix is created and are not allowed to be changed. If a constraint conflicts with one of the user-defined relationships, no subMAR can be found.

#### S4.2 Subpoplar, the sum rule algorithm

Here we describe our generalized sum rule algorithm Subpoplar, which is based on two key constraints: first, in a valid, complete clone tree all subclones must have a single parent, and second, the frequency of a subclone must be greater than or equal to the frequency sum of its children. Furthermore, we analyze Subpoplar’s runtime; *K* is the number of subclones including the germline and *N* is the number of samples.

Before Subpoplar starts, the possible parent matrix *τ* ∈ {0, 1}^*K×K*^ is created, following Definition 1 of a possible parent:

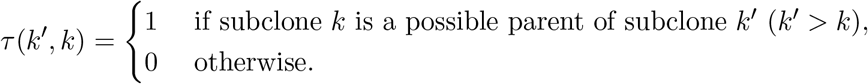

In addition, the vector *ψ*, storing the definite parent for each subclone, is created and initialized with *ψ*(*k*) = −1 for each subclone *k*. Also, a frequency matrix 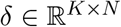 is created, which indicates the subclonal frequencies that subclones have available to become parents of other subclones without violating the sum constraint. It is initialized with the values of the subclonal frequency matrix *ϕ*. Creating *τ, ψ* and *δ* takes 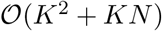 time.

Subpoplar processes the subclones in decreasing order of their average frequencies. The version of this algorithm working with subclonal CNAs is shown in Algorithms 5 to 8. For each subclone *k*, it is checked whether it can be a child of all its possible parents, hence whether its frequency is lower than or equal to the available frequencies of its possible parents in all samples (see Algorithm 5, lines 1-10). If subclone *k* has only one possible parent *k*, k* is made a definite child (see Algorithm 5, lines 11-17). This process involves decreasing the available frequency *δ*(*k*,n*) by *ϕ*(*k, n*) for each sample *n* (see Algorithm 6, lines 1-4). Furthermore, it is checked whether other possible children of subclone *k\** can remain its possible children or whether now after updating the available frequency *δ*(*k*, n*), they would violate the generalized sum rule if they became definite children (see Algorithm 6, lines 5-22). If they led to a violation, they are removed from the list of possible children. If they were already processed in a previous round of the algorithm and have now without *k\** only one possible parent left, the complete child updating process is performed recursively. At the end of each such process, the relationship *Z*(*k, k′*) is updated (see Algorithm 7). In basic mode of SubMARine, every update of an ancestral relationship also leads to a propagation of the partial tree rule (see Equation 1 and Algorithm 7, lines 33-35). In extended mode, in addition to the partial tree rule, SSM phases (see Algorithm 7, lines 20-21) and absent relationships (see Algorithm 7, lines 22-23 and Algorithm 8) are also propagated to satisfy the equivalence and lost allele constraints. At the end of Subpoplar, if a subclone does not have a definite parent yet, the lowest common ancestor of all its possible parents is made its ancestor to use all information present in the data to eventually propagate further relationships (see Algorithm 5, lines 18-25).

No matter whether Subpoplar was called from the basic or extended version of SubMARine, without children, relationship and SSM phasing updates, Subpoplar (Algorithm 5) needs 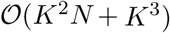 time because for each subclone and all of its possible parents, the frequencies in all samples have to be compared, and furthermore, all descendants of all possible parents might have to be processed. Making a child the definite child of its definite parent *k\** (Algorithm 6) without considering relationship updates and recursive calls, takes 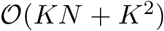 time because for each possible child of *k\** the frequencies in all samples have to be considered and eventually all possible descendants of *k\** have to be checked to be possible ancestors of the possible children. We now start to consider single relationship updates when updating children, differentiating the two possibly required values. If a possible child *k′* cannot be a descendant of *k\**, an absent relationship (*Z*(*k*,k′*) = 0) is created with Algorithm 7. Without considering further updates, updating to this absent relationship takes 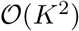 time because possible ancestors of *k\** have to be checked to have possible descendants that are possible parents of *k′*, and furthermore, the new relationship *Z*(*k*,k′*) can influence up to *K* relationships through the partial tree rule, which needs to be checked. Hence, making a child *k* the definite child of its definite parent *k\** and considering only absent relationship updates of this action, takes 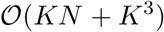 time, where one of the factors *K* from updating children is now superseded with *K*^2^. The second relationship update to consider when updating children is a positive relationship update (*Z*(*k*,k*) = 1). In basic mode, without SSM phasing and subclonal CNAs, and without further updates, this does not increase the asymptotic run time of updating children (see Algorithm 7). However, in extended mode, propagating SSM phasing and absent relationships to satisfy the equivalence and lost allele constraints takes an additional 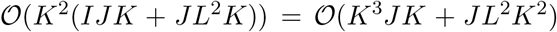 time (see Algorithm 7, line 20, and Algorithm 8, line 3 and equations mentioned therein), where I is the number of segments, *J* is the number of SSMs and *L* is the number of CNAs. Without further updates, making a child the definite child of its definite parent thus takes 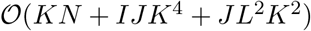, where analogously to the basic mode one factor *K* gets superseded by the complexity of the relationship update. Now, updating ancestral relationships can propagate further updates, yet, since each relationship is updated at most once, the number of total updates and hence relationship propagations is limited. Because there are only *K*^2^ relationships, the total runtime of the Subpoplar algorithm with all updates in basic mode is 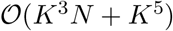 and in extended mode is 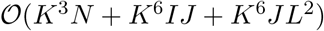.

##### Algorithm 5 Pseudocode of the Subpoplar algorithm

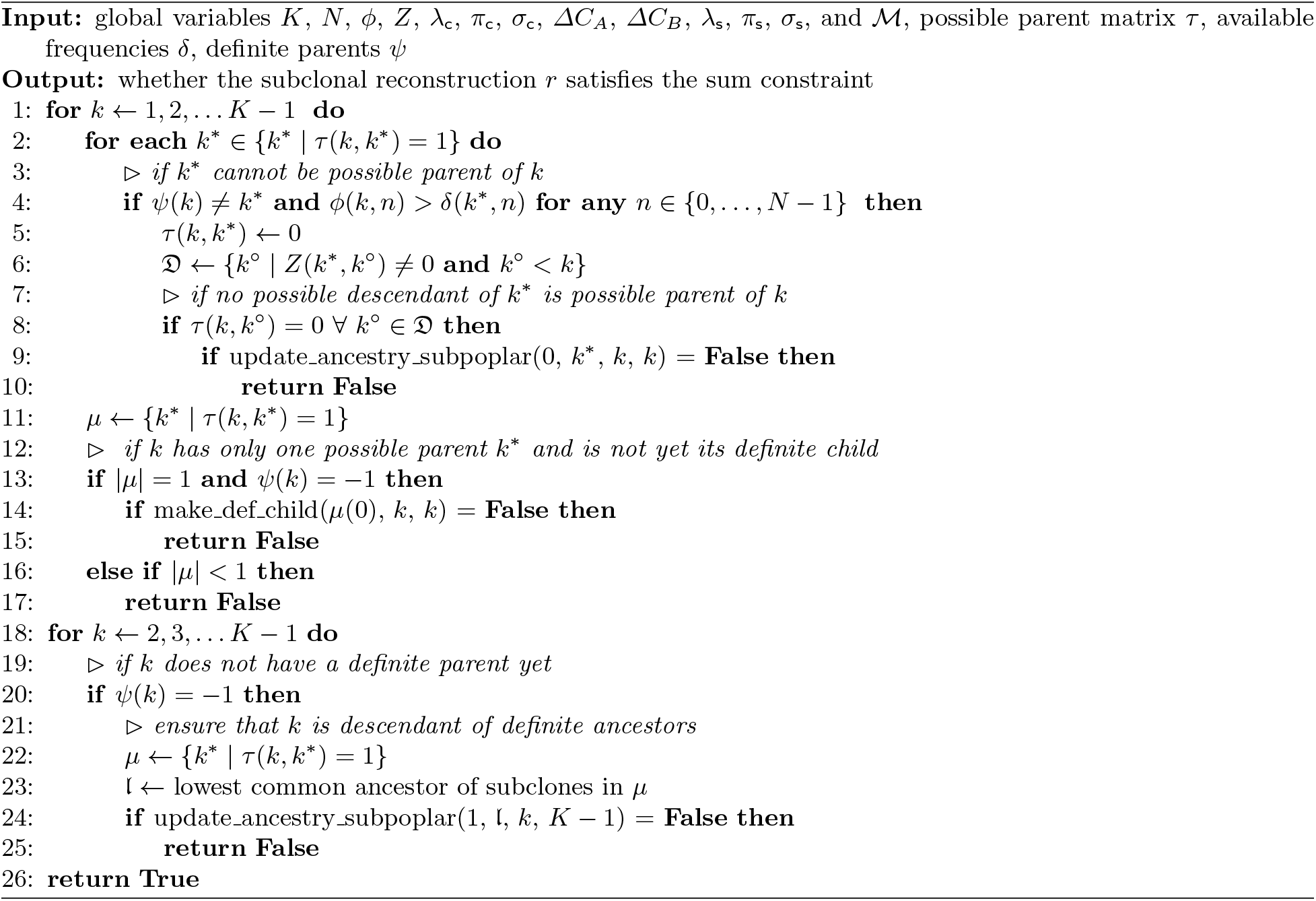

##### Algorithm 6 make_def_child(*k*, k, l*)

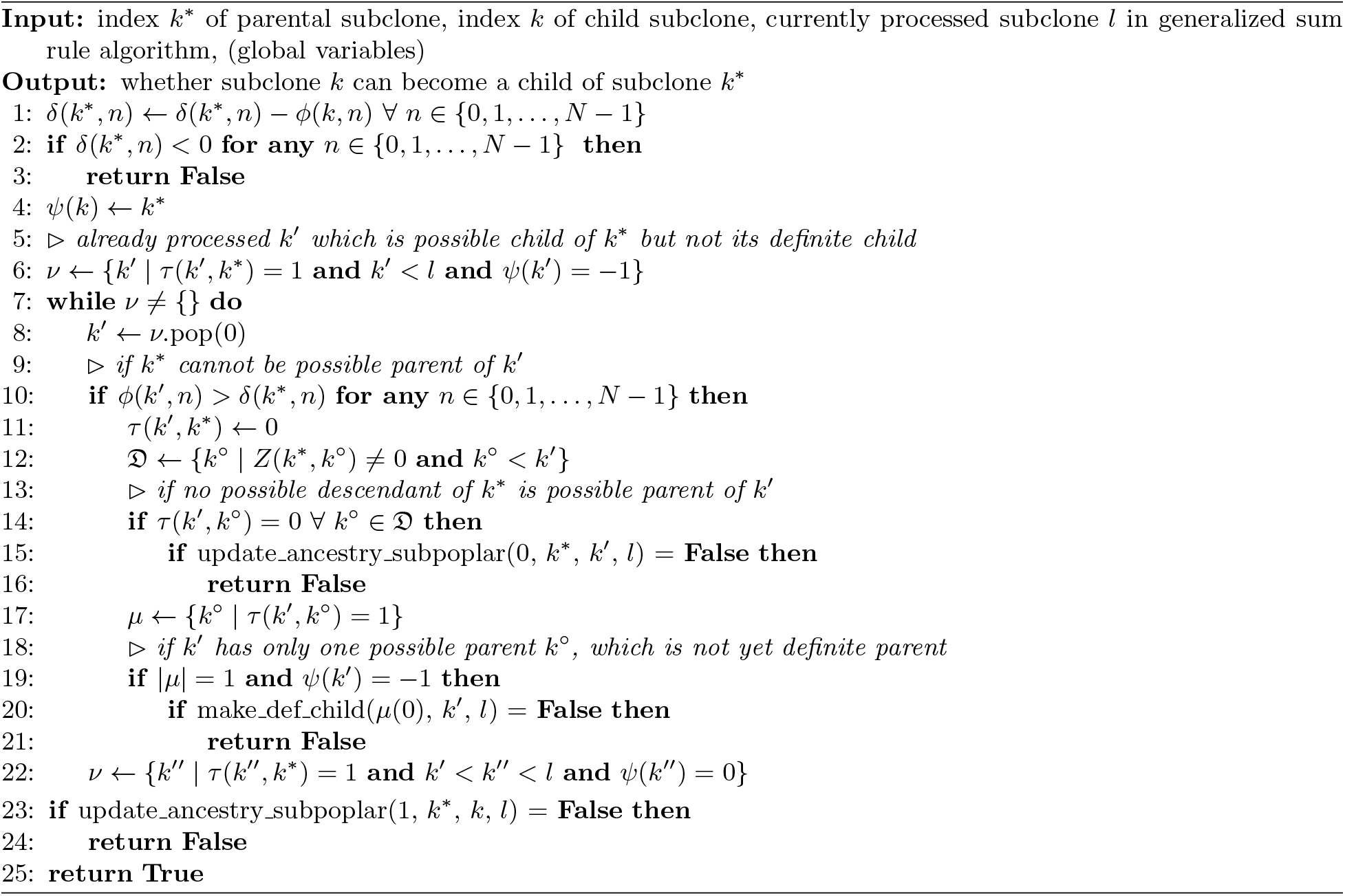

##### Algorithm 7 update_ancestry_subpoplar(*υ, k*, k, l*)

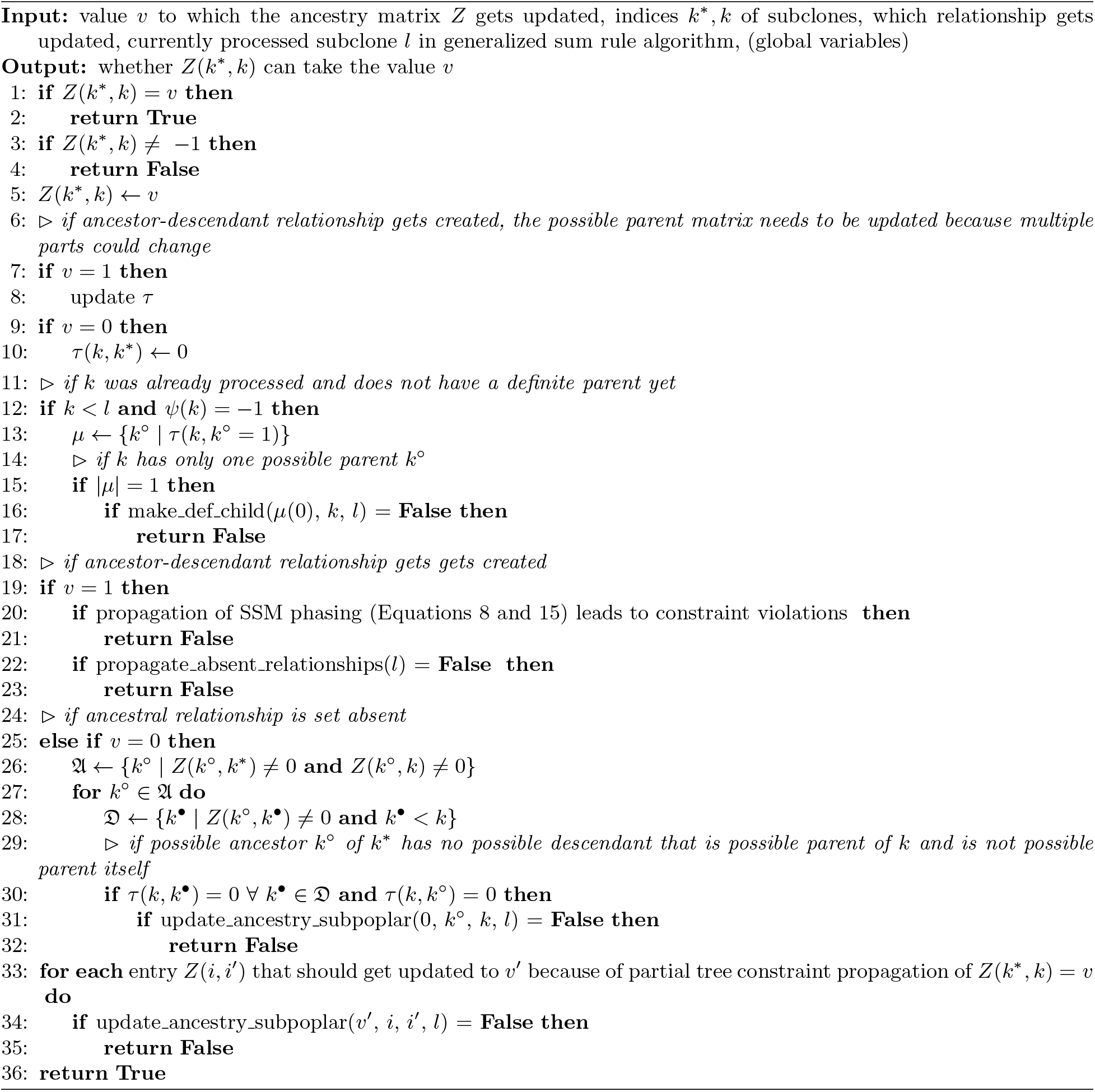

##### Algorithm 8 propagate_absent_relationships(l)

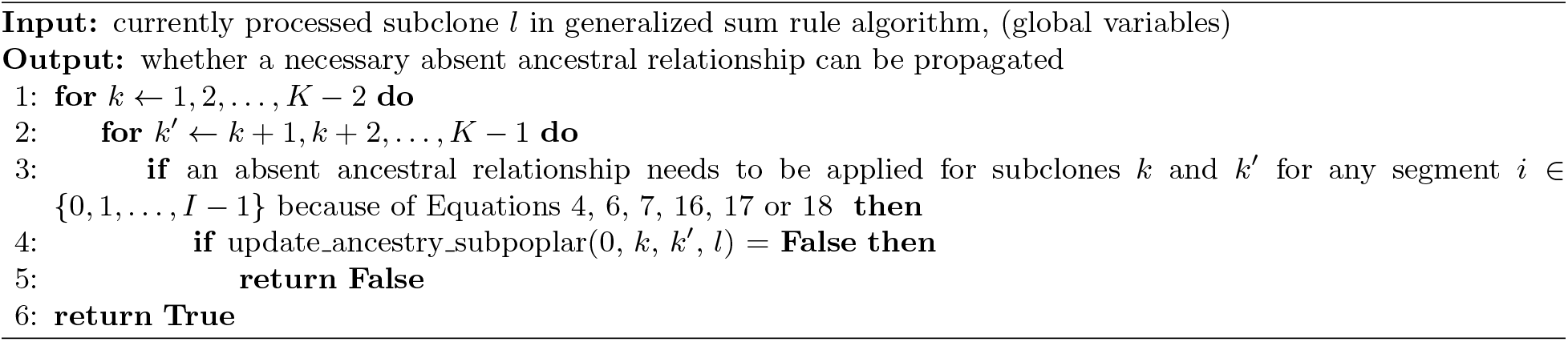

#### S4.3 Extending SubMARine to deal with noisy subclonal frequencies

The subclonal frequency matrix *ϕ* impacts directly the result of the generalized sum rule but not the result of any other inference rule (see Tables 1 and S1). The matrix is direct input to the crossing rule and Subpoplar. Indirectly, it influences setting the trivial relationships via the ordering of subclones. Because subclonal frequencies cannot be measured precisely from bulk cancer sequencing data but are inferred from noisy mutational frequencies, it is possible that no valid partial clone tree exists that satisfies the generalized sum constraint even though the infinite site assumption holds.

In order to deal with the issue of unprecise subclonal frequencies, we developed a noise-buffered version of SubMARine. We first describe how a minimum noise buffer uniform across subclones and samples is found in polynomial time and then how a subclone- and sample-specific buffer can be found.

In the original version of Subpoplar as described in Section S4.2, the algorithm checks for each subclone whether its subclonal frequencies are lower than or equal to the available frequencies of its possible parents. All possible parents for which this is not the case, are discarded. However, if the subclonal frequencies are inaccurate, it can happen that a subclone *k′* cannot be a child of any subclone *k*, not even of the germline. Hence, the generalized sum rule would require setting all entries *Z*(*k,k′*) = 0 but because *Z*(0,*k′*) = 1 as a consequence of the germline rule, no valid partial clone tree exists. To enable finding a partial clone tree also in these cases, we introduce the use of a noise buffer. This buffer is added to the parental frequencies, and leads to Subpoplar discarding possible parents only if the subclonal frequencies of a possible child are greater than the available frequencies of the possible parents plus this buffer *b*. This leads to the following changes in Subpoplar where we use *b*:

– Algorithm 5, line 4: **if** *ψ*(*k*) = *k\** **and** *ϕ*(*k, n*) > *δ*(*k*, n*) + *b* **for any** *n* ∈ {0,…, *N* − 1}
– Algorithm 6, line 2: **if** *δ*(*k*,n*) + *b* < 0 **for any** *n* ∈ {0,1,…, *N* − 1}
– Algorithm 6, line 10: **if** *ϕ*(*k′,n*) > *δ*(*k*,n*) + *b* **for any** *n* ∈ {0, 1,…, *N* − 1}

Also, we extend the crossing rule used in SubMARine to use the noise buffer *b*:

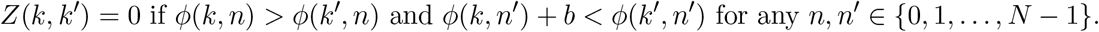

Whenever a subclone *k* has no possible parent because of the generalized sum constraint, we compute the minimum noise buffer *b* necessary so that *k* has at least one possible parent. Then, we start SubMARine again using the new buffer *b*. In the worst case, we have to increase the noise buffer once for each of the *K* subclones, still leading to a polynomial run time of SubMARine. However, the minimum noise buffer *b* computed for a subclone *k* is not necessarily the minimum noise buffer for the dataset (see Figure S15). Hence, once we find the valid subMAR for *b*, we use *b* as the starting point for a binary search to ensure finding this minimum buffer.

After applying the binary search, we have found the necessary minimum noise buffer uniform across all subclones and samples. However, while some subclones need this buffer in order for SubMARine to find a valid partial clone tree, other subclones might not need it to have at least one possible parent and might get more possible parents when using it. This leads to a subMAR with more uncertainty than necessary and to subMAR-completing trees varying in their data fit (see Figure S16). To prevent this from happening, SubMARine attempts to find in polynomial time, starting from the subMAR computed with the minimum uniform buffer, the subclone- and sample-specific noise buffer set and its corresponding subMAR, such that all completing clone trees have as little negative frequencies in their available frequency matrix *δ* as possible. For this purpose, for all subclones having multiple possible parents the available frequencies of all their possible parents get collected. The best possible subclone- and sample-specific buffer set is the one in which the lowest possible buffer is chosen for each subclone with multiple parents. Note that for all subclones that have only one possible parent, we choose the uniform buffer *b* as their specific buffer because even if a lower value is used, it has no influence on the subMAR-completing trees. If a valid subMAR exists for the best buffer set, SubMARine reports this subMAR and buffer set. Otherwise, SubMARine identifies the second best possible set and if a subMAR exists for it, reports this buffer set and subMAR. This second best set is the one where the subclone with the lowest second possible buffer chooses this buffer, and all other subclones with multiple possible parents choose their lowest possible buffer. If no valid subMAR exists for this second best set, SubMARine does not search for the third best one because in order to find it, all buffer combinations have to be considered which cannot be done in polynomial time anymore. Hence, SubMARine informs about the minimum uniform noise buffer and reports it along with the corresponding subMAR.

**Fig. S15.**
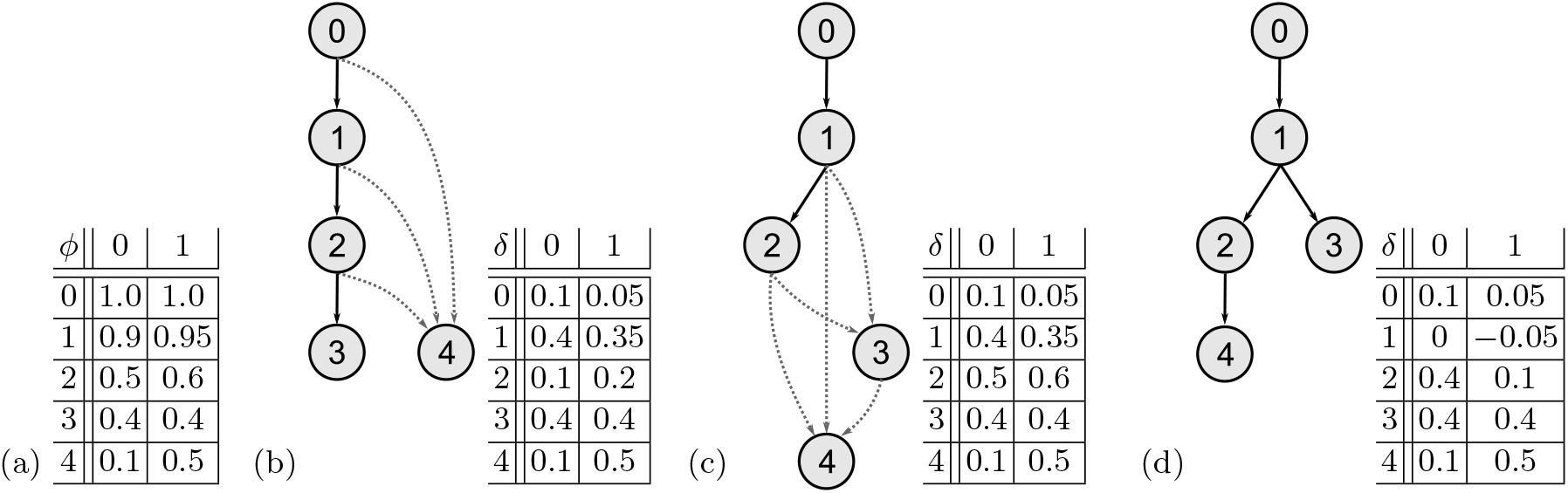
Example in which the minimum noise buffer for a subclone is not the minimum uniform noise buffer for the dataset and a binary search is necessary. (a) Subclonal frequency matrix *ϕ* for the germline and four subclones across two samples. (b) Temporary partial clone tree after applying Subpoplar up to subclone 3. The next step is to check for subclone 4 whether some of its three possible parents can be discarded because they do not have enough available frequencies in the matrix *δ* to become its definite parent. Since all have to be discarded, a noise buffer needs to be introduced to find a valid subMAR. The minimum noise buffer for subclone 4 to have at least one possible parent is 0.15. Then it could become a child of subclone 1. (c) Valid subMAR if SubMARine is run with a noise buffer of 0.15. Here, subclone 3 has two possible parents and subclone 4 has three. However, a lower noise buffer exists for this dataset. (d) Valid subMAR with a noise buffer of 0.05, identified through a binary search starting from 0.15. All subclonal relationships are defined.

**Fig. S16.**
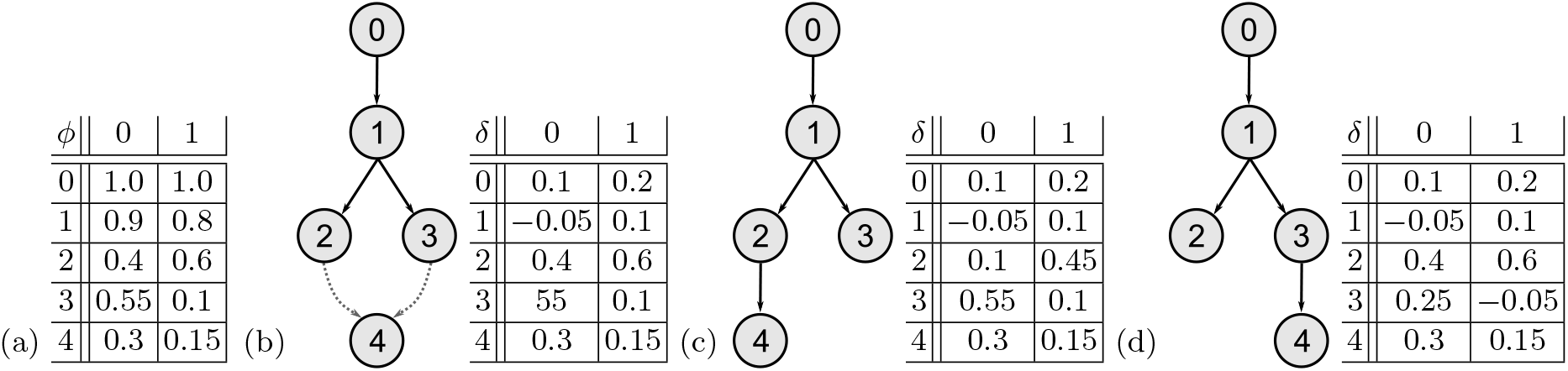
Example in which a uniform noise buffer leads to subMAR-completing trees with different data fit. (a) Subclonal frequency matrix *ϕ* for the germline and four subclones across two samples. (b) Valid subMAR and available frequency matrix *δ* when applying SubMARine with the lowest uniform noise buffer of 0.05 for this dataset. (c) SubMAR-completing tree where subclone 4 is a child of subclone 2. The available frequencies of subclone 2 stay positive. (d) SubMAR-completing tree where subclone 4 is a child of subclone 3. However, in order to become a child, the noise buffer is required. Thus, subclone 3 has a negative available frequency in sample 1. Hence, this tree fits the data worse than the tree in (c).

Instead of comparing all noise buffer combinations to find the best possible subclone- and sample-specific noise buffer set, we offer a different approach. Starting from the reported subMAR and using the uniform noise buffer, SubMARine can apply a depth-first search to find all completing clone trees that have as little negative frequencies in their available frequency matrix *δ* as possible and constructs their MAR. Whenever a clone tree *t* with an associated available frequency matrix *δ* is complete, SubMARine computes the amount of negative available frequency 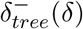 of this tree:

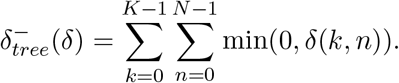

If this frequency is higher than the one of the previous completed tree or trees, they are discarded and the new clone tree t is kept. SubMARine computes the negative available frequency while completing a clone tree, thus can discard a partial clone tree as soon as its frequency gets smaller than the so far best one. When all trees got enumerated, SubMARine builds the MAR for the trees having as little negative available frequencies as possible and reports it along with this frequency and the used subclone- and sample-specific buffer set. Note that instead of keeping enumerated complete clone trees in memory, SubMARine stores in *K × K* matrix which subclonal relationships have been set to present and/or absent in the already enumerated trees. Whenever a tree with a better negative available frequency is found, the relationships in this matrix are set in accordance to this new tree.

#### S4.4 SubMARine with SSMs and clonal CNAs

When in addition to the subclonal frequency matrix *ϕ*, also SSMs and clonal CNAs are given, SubMARine can be used without any changes to build a subMAR, which describes the set of clone trees of all valid clone trees fitting this setting. The lost allele constraint, which needs to be satisfies when CNAs are given and usually requires knowing the subclone and allele assignment of SSMs, does not have to be propagated. The reason is that as long as only one allele of a segment is deleted by a clonal CNA, SSMs in this segment can be phased to the other allele, which is not affected by any other CNA. Only if there are clonal losses on both alleles in one segment that also contains SSMs, no valid clone tree exists for this input. We test for this special scenario with a verification step prior to SubMARine that requires the parental allele assignment of the clonal CNAs, and the segment assignment of the CNAs and SSMs. The verification adds the term 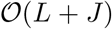 to the runtime of SubMARine, where *L* is the number of CNAs and *J* is the number of SSMs.

#### S4.5 Upper bound on the size of MAR- and subMAR-completing clone trees

The MAR and the subMAR are a summary of the set of clone trees that complete them and that are all valid. Counting the number of all valid clone trees given a basic clone tree reconstruction problems was shown to be #P-complete [36]. Hence, we derive an upper bound in polynomial time by considering the *possible parents* of each subclone. A possible parent of a subclone includes all ancestral subclones of which this subclone is a child or could become a child. Note that a subclone could also have a single possible parent. Without application of the Subpoplar algorithm (see Section S4.2), a possible parent is formally defined as follows:

#### Definition 1 (Possible parent).

*Subclone *k* is a possible parent of subclone *k′* if*

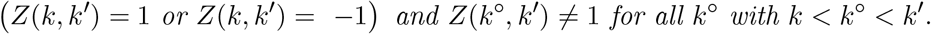

If the Subpoplar algorithm was applied, possible parents of a subclone *k′* are indicated in row *τ*(*k′*) of the possible parent matrix *τ*, which is returned by the algorithm.

The number of trees in the summary set of clone trees can easily be computed as follows:

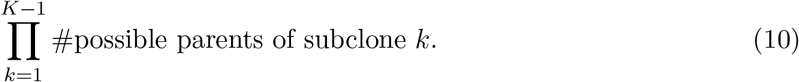

However, because this set can contain clone trees that do not complete the MAR or subMAR (see Figure S17), its size is an upper bound.

**Fig. S17.**
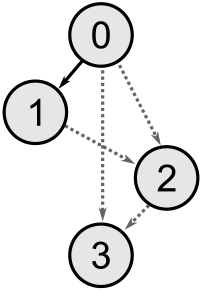
Example of a partial clone tree that describes a clone tree not completing it. Subclones 2 and 3 have two possible parents and subclone 1 cannot be an ancestor of subclone 3. When choosing subclone 2 as parent for subclone 3, and subclone 1 as parent for subclone 2, subclone 1 has to be an ancestor of subclone 3 because of the transitivity property of the tree constraint. Then however, this constructed tree does not complete the clone tree anymore.

### S5 Details on extended SubMARine

#### S5.1 Computation of implied copy numbers and VAFs

The data fit to the experimentally-derived average copy numbers of the segments and the VAFs of the SSMs depends on the average copy numbers and VAFs implied by the clone tree and mutation assignments. Here, we describe how these can be computed and which role the ancestral relationships between subclones play.

The implied average allele-specific copy number for allele *A* in segment *i* of sample *n* can be computed as:

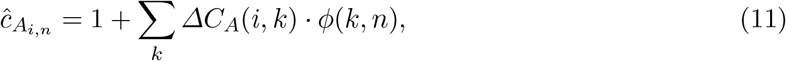

where 1 is the normal copy number of the allele and *ΔC_A_*(*i, k*) is the copy number change of allele *A* in segment *i* of subclone *k* as defined in Section S2. 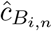 can be expressed analogously. Note that 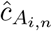 and 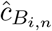 do not depend on the ancestral relationships between subclones.

The implied VAF of SSM *j* in sample *n* can be computed as:

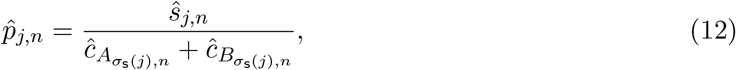

where 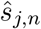 is the mutant copy number of the SSM and is computed as follows:

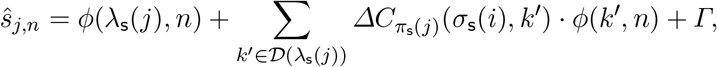

where the function 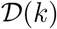 returns all descendants of subclone *k*:

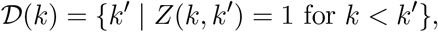

and *Γ* is defined as follows:

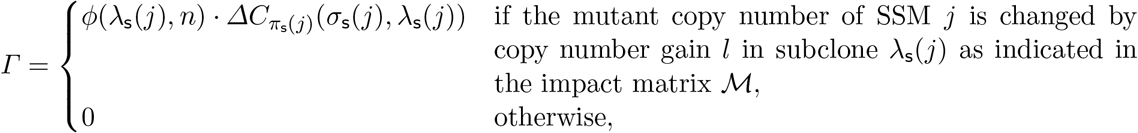

where λ_s_(*j*), *π*_s_(*j*) and *σ*_s_(*j*) are the subclone, parental allele and segment assignments of SSM *j* as defined in Section S2, and 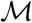 is the impact matrix as defined in Section 4. Because the descendants of subclone λ_s_(*j*) have an influence on the computation of the VAF, the ancestral relationships between subclones matters.

#### S5.2 Same data fit despite different impact matrices

In order for two clone trees *c* and *c′* with the same subclonal frequencies and mutation assignments to infer the same data fit although they differ in their impact matrices, one of the following conditions has to hold:

1. two subclones with the same copy number change in the same segment on the same allele have to have exactly the same subclonal frequencies in all samples,
2. two sets of subclones with different subclonal frequencies have to result in exactly the same copy number changes in the same segment on the same allele in all samples. Figure S18 shows two examples of these conditions.

#### S5.3 Equivalence constraints

The segment, parental allele and subclonal assignments of CNAs and SSMs are stored in the vectors *σ*_c_, *π*_c_, λ_c_ and *σ*_s_, *π*_s_, λ_s_, respectively, as was defined in Section S2.

If a CNA *l* changes the mutant copy number of an SSM *j* and both are assigned to different subclones, then the CNA’s subclone has to be descendant of the SSM’s one:

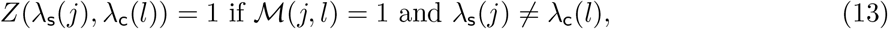

**Fig. S18.**
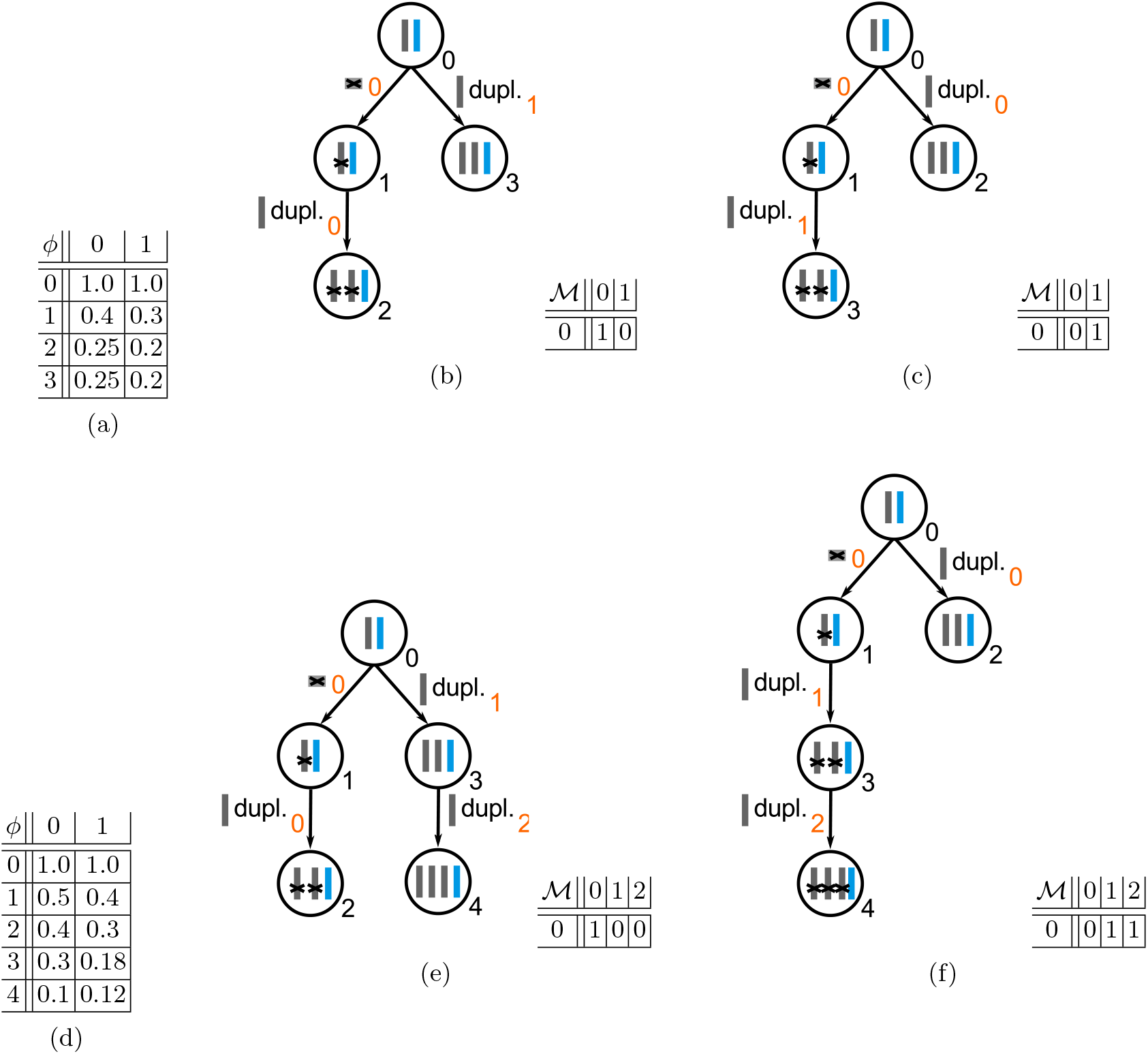
Two examples with different clone trees that have the same subclonal frequencies and mutation assignments but different impact matrices and still infer the same data fit. (a)–(c) Subclones 2 and 3 have the same subclonal frequencies across both samples and contain both a copy number gain. Thus, they influence SSM 0 in the same way which leads to the same VAF. (d)–(f) Whether the mutant copy number of SSM 0 is changed only by copy number change 0 in subclone 2 or by both copy number changes 1 and 2 in subclones 3 and 4 leads to the same VAF (see Equation 12).

Furthermore, the SSM *j* needs to be phased to the same allele as the CNA *l*:

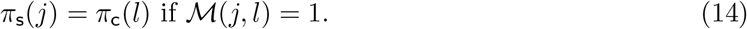

If the mutant copy number of an SSM *j* should not be changed by a CNA *l* but the CNA is assigned to a descendant subclone in the same segment, the SSM needs to be phased to the opposite allele:

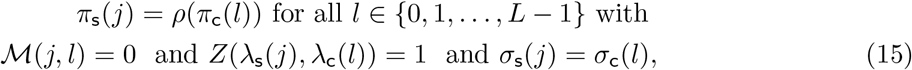

where the function *ρ*(*α*) returns the opposite allele:

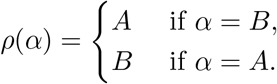

If the phase of an SSM cannot be adapted in order to avoid the unwanted influence of a CNA, the ancestral relationship between the subclone with the SSM and the one with the CNA needs to be absent. There exist three cases where this occurs:

1. If subclone *k* has an SSM *j* and subclone *k′* has a CNA *l* that are both assigned to the same segment and allele but the CNA should not change the copy number of SSM *j*, the ancestral relationship between the two subclones has to be absent:

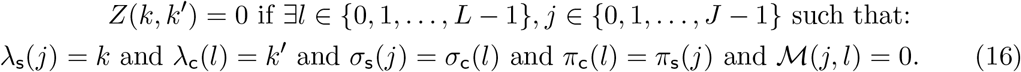
2. If subclone *k* has an SSM *j*, subclone *k′* has a CNA *l* in the same segment and either subclone *k′* or its descendant *k″* have another CNAs *l′* in the same segment on the other allele than *l*, subclone *k* cannot be an ancestor of subclone *k′* if the copy number of SSM *j* should not be changed by the two CNAs:

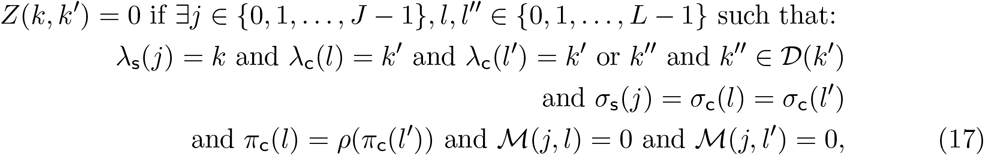

where the function 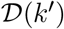 returns all descendants of subclone *k′* and is defined in Section S5.1.
3. If subclone *k′* has a CNA *l* on one allele and is the descendant of a subclone that lost all copies of the other allele in the same segment, it cannot be the descendant of subclone *k* that has an SSM *j* in the same segment whose mutant copy number should not be changed by the CNA *l*:

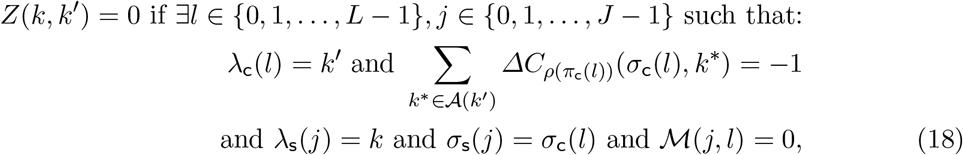

where the function 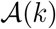 returns all ancestors of subclone *k* and was defined in Section S2.

#### S5.4 Monotonicity restriction

To ensure that the input provided to SubMARine has only copy number changes in one direction per segment and allele, it has to satisfy the following two *monotonicity constraints*:

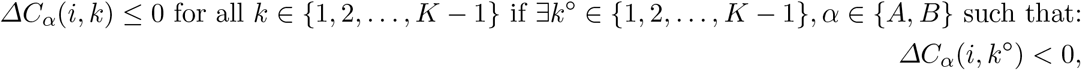

and

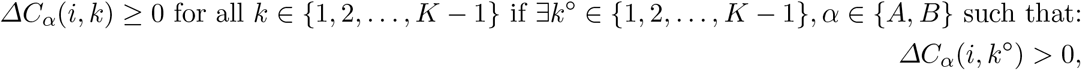

where *ΔC_A_* and *ΔC_B_* describe the copy number change per allele, segment and subclone and are defined in Section S2.

Without the input satisfying the monotonicity constraints, SubMARine could not guarantee that all defined ancestral relationships and SSM phases specified by inference rule propagation in its extended subMAR have the same value in the corresponding extended MAR (see Figure S19).

#### S5.5 Example of SubMARine in extended mode

This section contains an example how SubMARine works. For a detailed description with a runtime analysis see Section S5.6.

Given the subclonal frequency matrix *ϕ*, the impact matrix M, all CNA information, and the SSM assignment to segments and subclones as input (see Figure S20c-e), SubMARine in extended mode builds the valid partial clone tree in Figure S20a in the following order.

Because the monotonicity restriction holds on the CNAs, the preprocessing phase applies the germline rule and sets all trivial relationships *Z*(*k,k′*) = 0 with *k′ ≤ k*.

Then the main phase starts. First, those of the equivalence and lost allele rules are propagated that lead to 1’s in the ancestry matrix *Z* or that update SSM phasing. Whenever a relationship is updated, the partial tree rule is applied as well. Because CNA 0 of subclone 5 changes the mutant copy numbers of SSMs 0 and 2 of subclones 1 and 4 (Figure S5.6d,e), the SSMs are phased to the same allele as the CNA (equivalence rule based on Equation 14) and subclone 5 has to be a descendant of subclones 1 and 4 (equivalence rule based on Equation 13). In order to satisfy the single parent property of the partial tree constraint (Equation 1), subclone 1 needs to be an ancestor of subclone 4. CNA 1 of subclone 6 influences the mutant copy numbers of SSMs 1 and 5 of subclones 1 and 6 (Figure S5.6d,e). Hence, both SSMs are phased to the same allele as the CNA (equivalence based on Equation 14) and subclone 1 has to be an ancestor of subclone 6 (equivalence rule based on Equation 13). Because the mutant copy numbers of SSMs 1 and 3 should not be influenced by CNA 0 (Figure S5.6d), they get phased to the other allele (equivalence rule based on Equation 15). (SSM 1 was phased to this other allele already because of CNA 1 and Equation 14.) SSM 4 appears after the loss of allele B (Figure S5.6d,e), hence it has to be phased to allele A (lost allele rule based on Equation 8).

Second, those of the equivalence and lost allele rules that lead to 0’s in *Z*, and the crossing rule (Equation 9), which follows from the generalized sum constraint (Equation 3) and also leads to 0’s in *Z*, are propagated together with the partial tree constraint. Since SSM 4 is phased to the same allele as CNA 1 but its mutant copy number is not influenced by it (Figure S5.6d), subclone 5 of the SSM cannot be an ancestor of subclone 6 of the CNA (equivalence rule based on Equation 16). The same reasoning holds for SSM 0 and CNA 2, which is why subclone 1 of the SSM cannot be an ancestor of subclone 3 of the CNA. The transitivity property of the partial tree constraint leads to subclone 3 not able to be an ancestor of subclones 4, 5 and 6. In addition to this, subclone 3 could not be an ancestor of subclone 4 because of the lost allele rule based on Equation 6. The crossing rule forbids subclone 2 to be a descendant of subclone 1 (Figure S5.6c). Thus, subclone 2 cannot be an ancestor of subclones 4, 5, and 6 (transitivity property of the partial tree constraint).

**Fig. S19.**
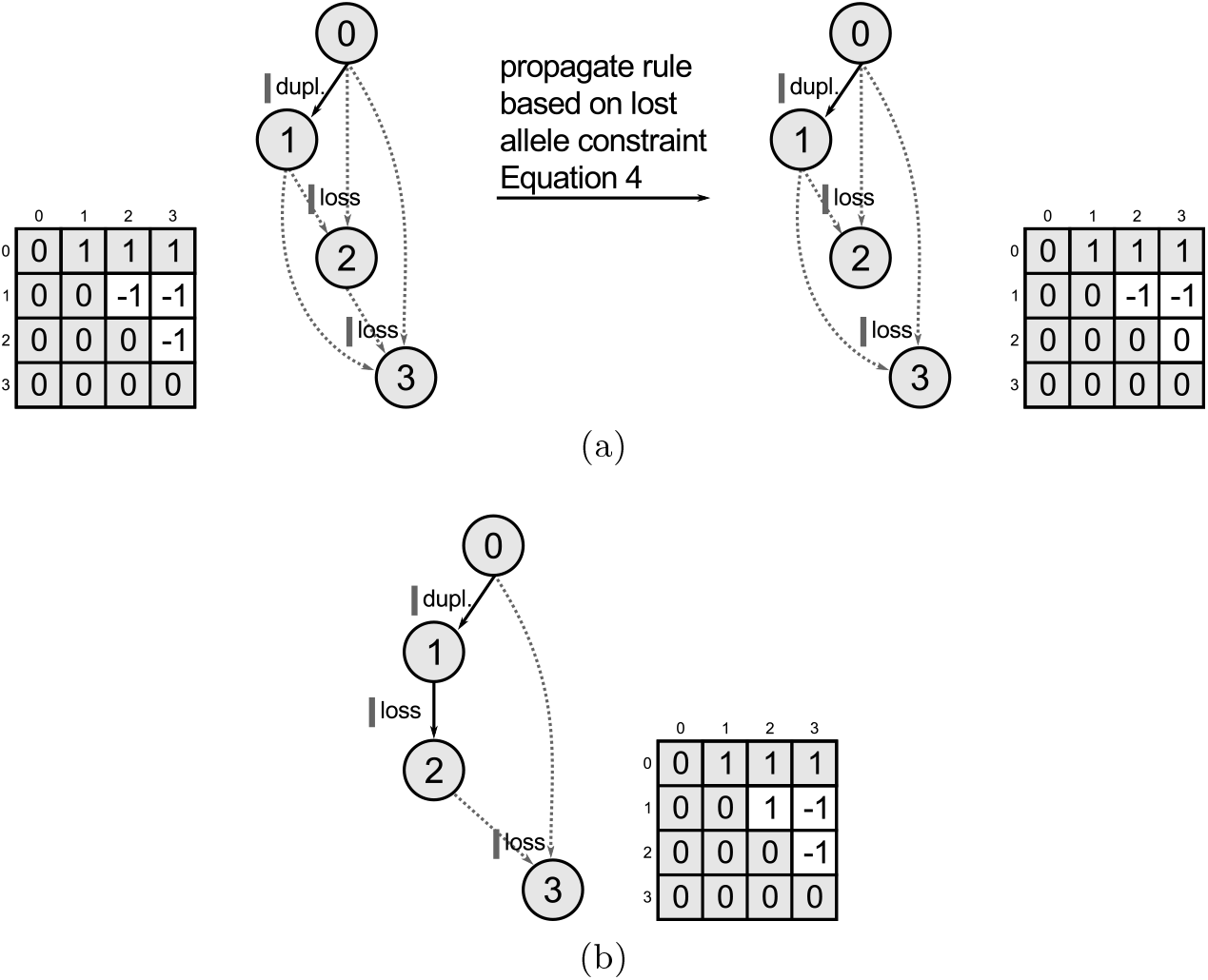
Example why extended SubMARine cannot guarantee that defined ancestral relationships in a subMAR have the same value in the corresponding MAR when the monotonicity constraint does not hold. (a) Partial clone tree with ancestry matrix *Z* before and after inference rule propagation. Subclonal frequencies are not shown because they are not relevant for this example. Gray entries in ancestry matrix *Z* are trivially a consequence of the ordering of the subclones and the generalized sum rule. Before propagating rules based on the lost allele constraint, subclone 3 can be a descendant of all other subclones. Due to a rule based on Equation 4, subclone 2 is not allowed to be an ancestor of subclone 3 and the corresponding undefined value in the ancestry matrix has to be updated to Z(2, 3) = 0. After this update, no more rules affect the undefined values of entry Z(1, 2) or Z(1, 3), hence, SubMARine terminates. (b) If now, the undefined relationship between subclones 1 and 2 was set to a definite one, making subclone 2 a descendant of subclone 1, then because the allele lost in subclone 2 was duplicated in subclone 1 and hence not all copies were lost, subclone 2 can be a possible ancestor of subclone 3 again. Thus, there exists a valid and equivalent clone tree that does not complete the subMAR and as a consequence, a defined entry in the subMAR would be undefined in the MAR.

**Fig. S20.**
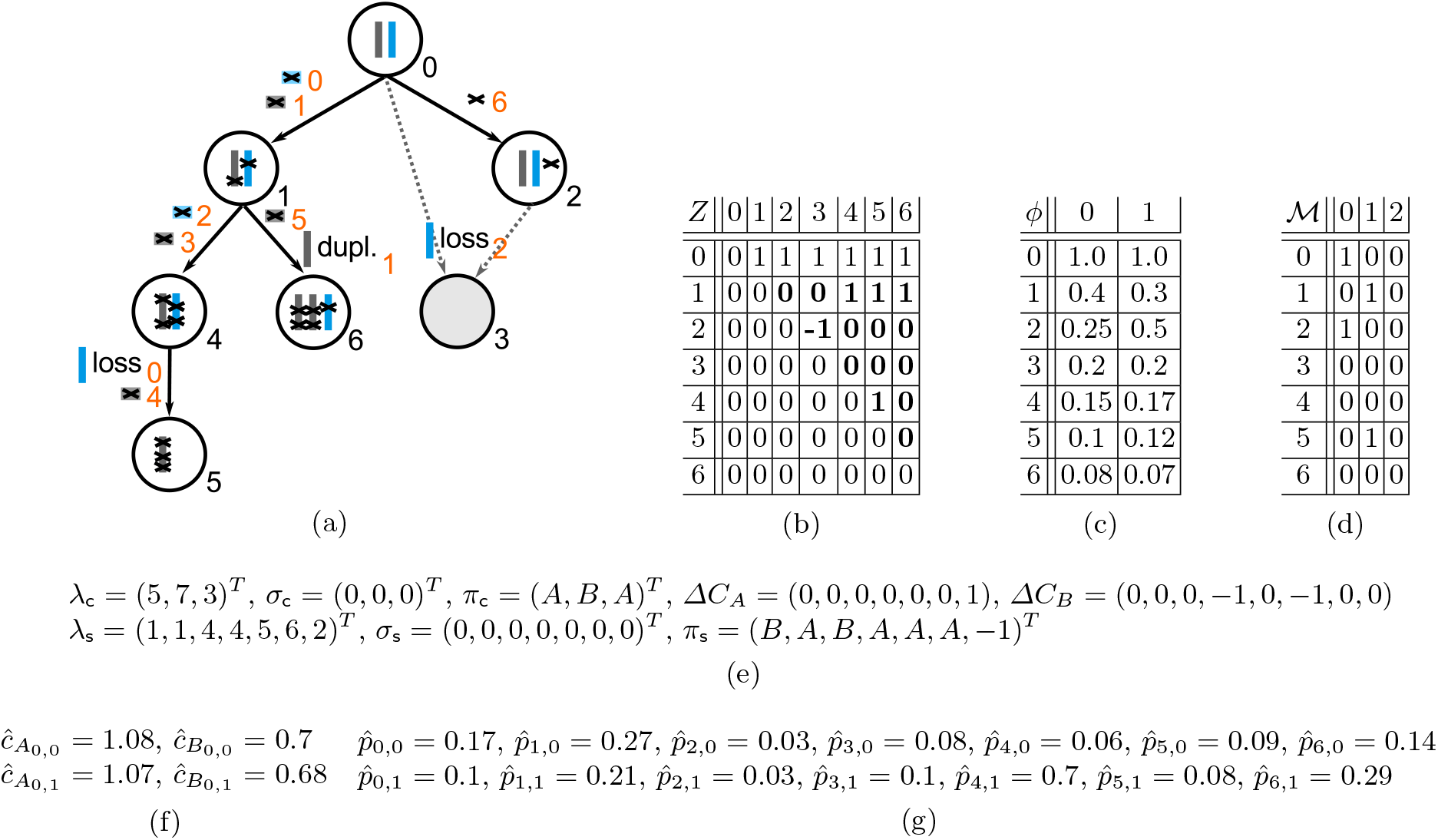
Valid partial clone tree (a) drawn as partial tree and (b) represented as ancestry matrix *Z* with (c) subclonal frequency matrix *ϕ*, (d) impact matrix 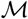 showing influence of CNAs on SSMs, (e) subclone, segment and parental allele assignments for CNAs and SSMs and type of copy number change for CNAs, (f) inferred average copy numbers, and (g) inferred VAFs. This partial clone tree consists of the germline (at the top of (a) with black index 0) and six subclones. We assume that only one segment is given. Allele *A* is duplicated in subclone 6, allele *B* gets lost in subclones 3 and 5. Four SSMs are phased to allele A, two are phased to allele B and one is unphased. (Indices of mutations are shown with orange numbers.) Every subclone but subclone 3 has a single possible parent. The two possible parents of subclone 3 are the germline and subclone 2. Thus, the genotype of subclone 3 cannot be unambiguously determined.

Third, the generalized sum rule is propagated with Subpoplar. Per default, the germline is the parent of subclone 1. Because subclone 2 cannot be a descendant of subclone 1, the germline is its parent as well. The subclonal frequencies allow subclone 3 to either be a child of the germline or of subclone 2, hence it has two possible parents. Subclones 4, 5 and 6 have only one possible parent left, hence no relationships have to be updated and no inference rule be propagated. Thus, SubMARine terminates and outputs the valid partial clone tree, represented by *Z*, the SSM phasing vector *π*_s_ and the possible parent matrix *τ* (not shown here).

#### S5.6 SubMARine in extended mode

We now describe in detail the extended mode of SubMARine, which approximates the extended maximally-constrained ancestral reconstruction problem, and analyze its runtime. *K* is the number of subclones including the germline, *N* is the number of samples, *I* is the number of segments, *J* is the number of SSMs and *L* is the number of CNAs.

The extended version of SubMARine (Algorithms 9-11) takes the subclonal frequencies *ϕ, L* CNAs with segment, subclonal and phase assignment, *σ*_c_, λ_c_ and *π*_c_, respectively, the direction and magnitude of copy number changes *ΔC_A_* and *ΔC_B_* for each allele derived from the CNAs, *J* SSMs with segment and subclonal assignment, *σ*_s_ and λ_s_, respectively, and the impact matrix 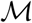 of an equivalent clone tree reconstruction problem t as input (see Figures S5 and S20). (More information on the notation of mutation assignments can be found in Section S2.) As in basic mode (see Section S4.1), the subclones are sorted in decreasing order of their subclonal frequencies. Extended SubMARine starts by creating an ancestry matrix *Z* in which all relationships are initially undefined 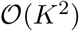 time, line 3 of Algorithm 9). Additionally, the phases of all SSMs are initialized in the vector *π*_s_ with the undefined value (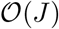 time, line 7). Then the monotonicity restriction is checked in 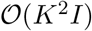 time (line 9). In the preprocessing phase, the germline rule is introduced (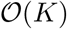 time lines 13 and 14), and trivial relationships are set as a consequence of the generalized sum rule and sorting of subclones, i.e. *Z*(*k,k′*) = 0 with *k′ ≤ k* (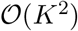 time, lines 15-17). Then the main phase starts and extended SubMARine takes care that SSMs are influenced by CNAs as indicated by the impact matrix 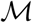 (line 20 of Algorithm 9 and Algorithm 10). First, it propagates Equation 13 and phases SSMs to the alleles of CNAs that impact them and creates ancestral-descendant relationships (*Z* (*k,k′*) = 1) between subclones (lines 1-13 of Algorithm 10). If no other ancestral relationships get propagated by the partial tree rule, which are checked after creating an ancestral-descendant relationship, this takes 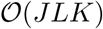 time. Because of the possible creation of ancestral-descendant relationships, SSMs not impacted by CNAs that are now in a descendant subclone must be phased to the other allele; this is done by propagating Equation 15 (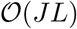 time, lines 15-21). After ensuring that the equivalence constraints are satisfied so far, the lost allele constraint needs to be checked. This is done by propagating Equation 8 and updating SSM phasing whenever an SSM could be phased to a lost allele otherwise (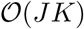 time, lines 23-29). In total, taking care that SSMs are influenced by CNAs as indicated by the impact matrix M and that the equivalence and lost allele constraints are satisfied takes 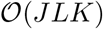 time when the partial tree rule does not lead to the propagation of further relationships. Now, to ensure that the equivalence and lost allele constraints are satisfied, absent ancestral relationships (*Z*(*k,k′*) = 0) are propagated (line 22 of Algorithm 9 and Algorithm 11). First, Equations 16, 18 and 17 of the equivalence constraint are propagated (lines 2-17 of Algorithm 11), which takes 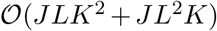 time if no other relationships get propagated because of the partial tree rule. Second, Equations 4, 6 and 7 of the lost allele constraint are propagated (lines 19-37), taking 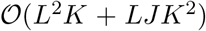 time with no relationship updates caused by the partial tree rule. Note that because of the monotonicity restriction, Equation 5 does not have to be considered. In total, propagating absent relationships with Algorithm 11 takes 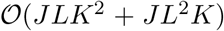 time without further updates. Afterwards extended SubMARine uses the crossing rule (Equation 9), which includes propagating the partial tree rule (lines 25 and 29 of Algorithm 9). This can be achieved in 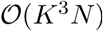 time when no other relationships are propagated by the tree rule and the crossing rule is implemented with the trick described in Section S4.1. Before considering the last step of extended SubMARine, which is propagating the generalized sum rule with the Subpoplar algorithm, we summarize extended SubMARine’s runtime so far. Without relationship updates caused by propagating the partial tree rule, it has a runtime of 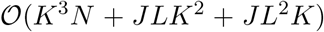. Because the ancestry matrix has only *K*^2^ ancestral relationships and each relationship is updated at most once, the total runtime of extended SubMARine so far when considering relationship updates of the partial tree rule is simply 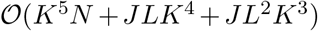. Finally, the generalized sum rule is propagated with the Subpoplar algorithm, which also takes care of the partial tree, the equivalence and the lost allele rules. In Section S4.2, we present this algorithm, which also creates and updates a possible parent matrix *τ*, indicating the possible parents for each subclone. Additionally, we derive its runtime of 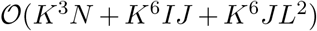, which already considers all possible relationship updates. Hence, the total runtime of extended SubMARine is 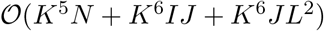.

Extended SubMARine converges when no ancestral relationship or SSM phases can be propagated anymore, which is after the Subpoplar algorithm finishes. Because only undefined relationships and SSM phases are updated and those are finite, it always converges. It returns an extended subMAR as result, which consists of the ancestry matrix *Z*, the SSM phasing vector *π*_s_ and the possible parent matrix *τ*.

It is possible for a user to define relationships for subclones and phases for SSMs. These relationships and phases are set after the initialization of *Z* and *π*_s_ (see Figure S5) and are not allowed to be changed. If a constraint conflicts with one of the user-defined relationships, no subMAR can be found.

Like the basic subMAR, the extended subMAR has three important properties for an extended clone tree reconstruction problem *t*: its defined ancestral relationships and SSM phases are a subset of those in the extended MAR, it is unique, and consequently, all valid and equivalent clone trees of t are completions of the extended subMAR. The reasoning for this follows the same argument as for the basic subMAR. Only undefined relationships and SSM phases are updated to defined ones and only when, given all other defined values, one of the two possible defined value causes a violation of a validity or equivalence constraint. Because the input data satisfies the monotonicity restriction, no updated value can be transformed back to the undefined value without violating a rule. Hence, the defined values are a subset of those in the extended subMAR. Even though the extended version of SubMARine works with more inference rules, i. e. those belonging to the lost allele and the equivalence constraints, no rule depends on an undefined value in order to update another undefined value. Thus, given a set of initially defined values, the order in which the inference rules are applied does not matter; the extended subMAR is unique.

**Algorithm 9.**
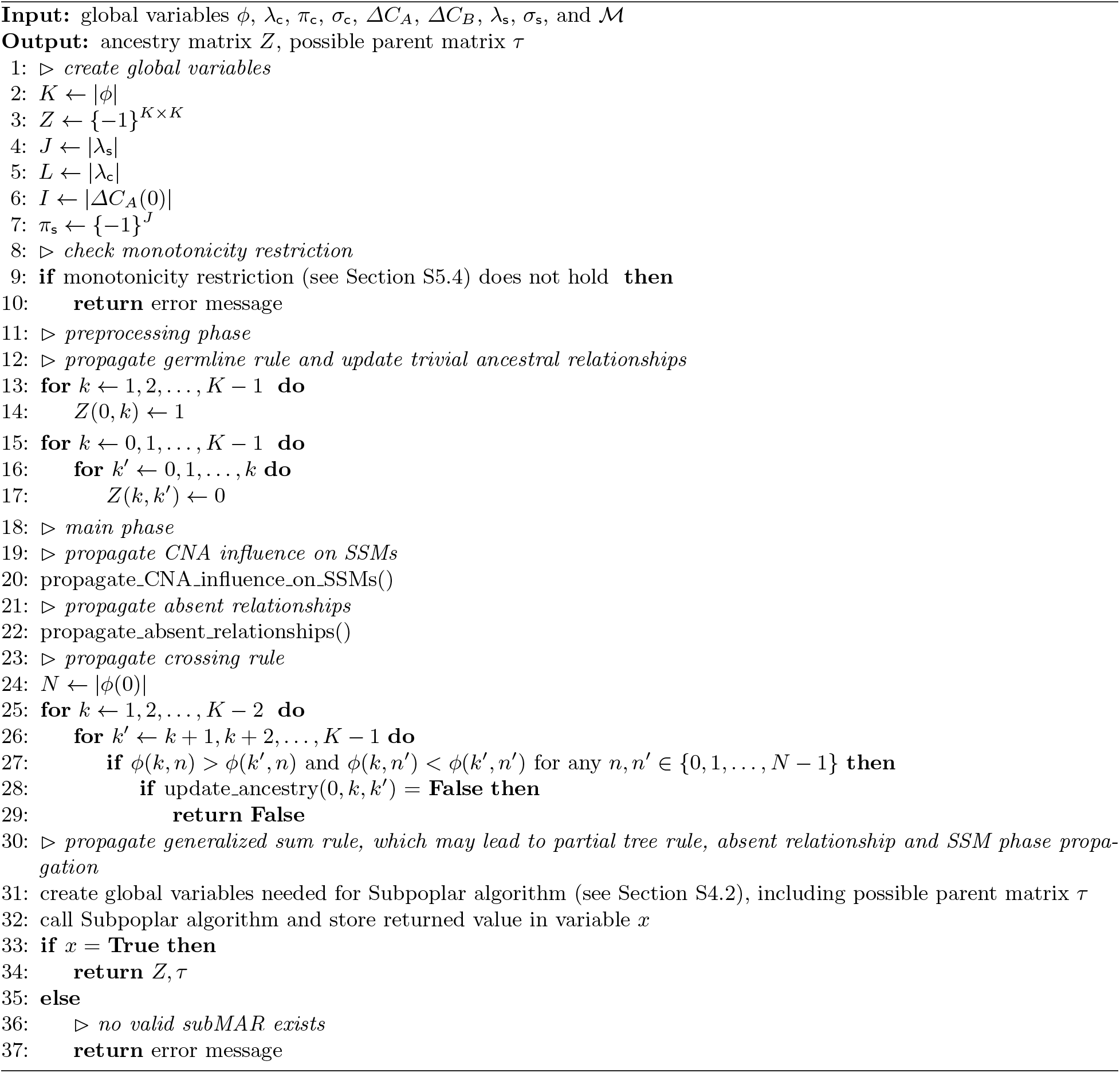
Pseudocode of the SubMARine algorithm in extended mode

**Algorithm 10.**
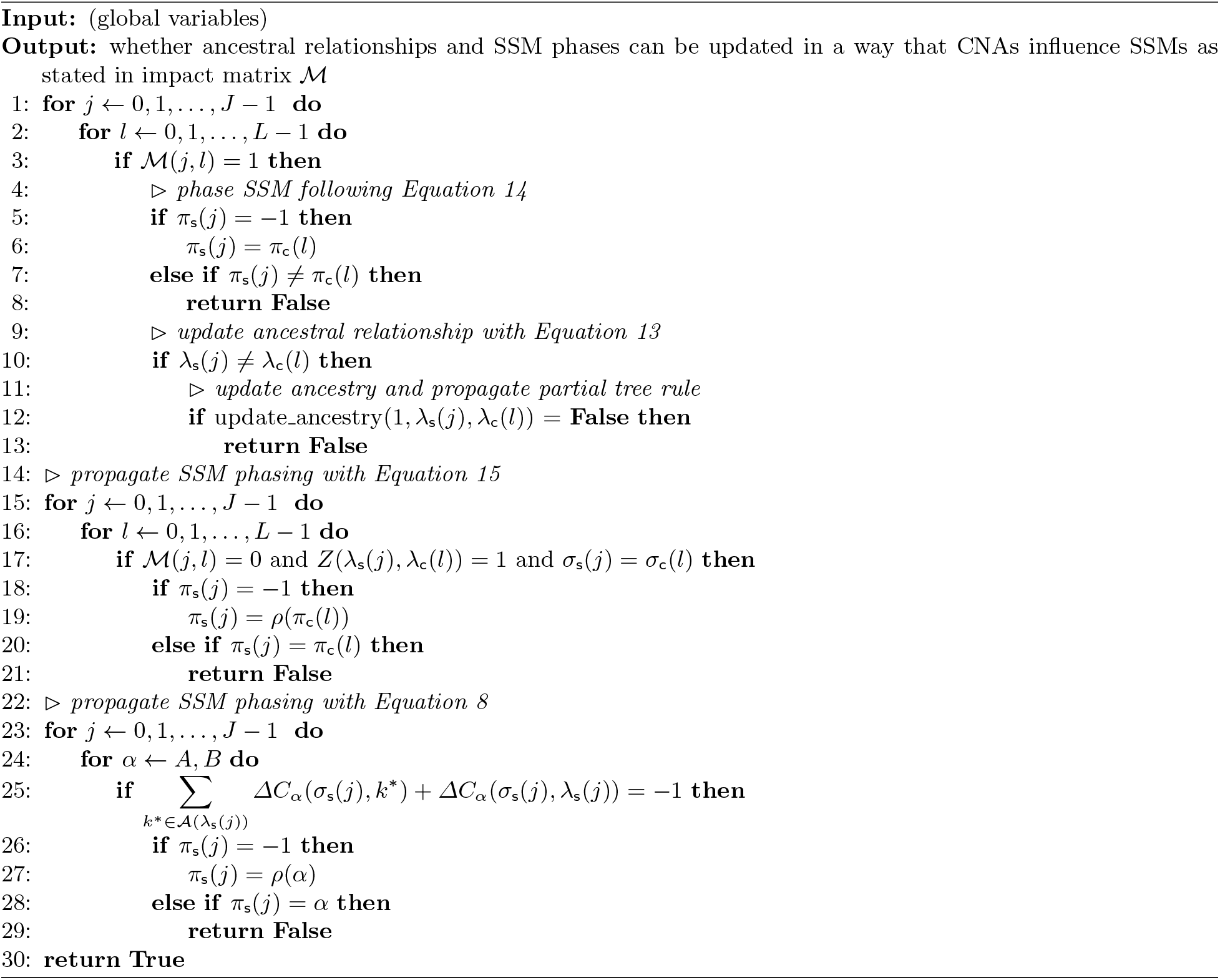
propagate_CNA_influence_on_SSMs()

**Algorithm 11.**
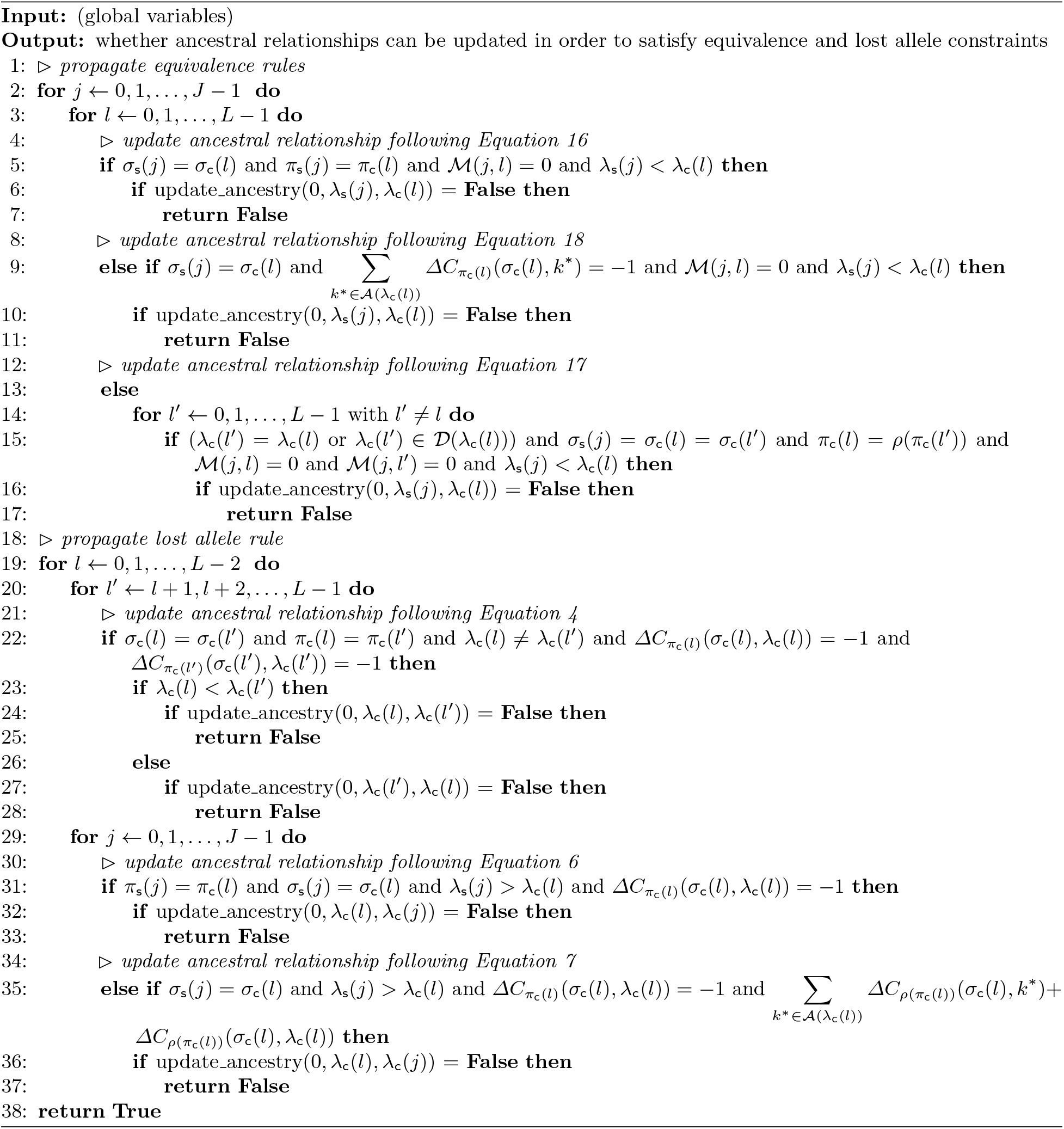
propagate_absent_relationships()

### S6 Details on results

#### S6.1 Simulating subclonal reconstructions

To simulate subclonal reconstructions, we first define parameters controlling the simulated data:

– *K*: number of subclones including the germline
– *N*: number of tumor samples
– *J*: number of SSMs
– *L*: number of CNAs
– *I*: number of genomic segments

We then generate simulated data using the following procedure:

1. Generate the tree structure. For each subclone *k*, with *k* ∈ {1, 2,…, *K* − 1}, sample a parent 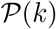. We extend the previous subclone (i.e., 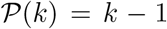) with probability *μ* = 0.75, and otherwise sample 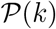 from the discrete Uniform(0, *k* − 1) distribution.
2. Generate the population frequencies *η*(*k, s*) for each population *k* in each tumor sample *s*, with *s* ∈ {0, 1,…, *N* − 1}. These values were sampled for each *s* as {*η*(0, *s*), *η*(1, *s*),…, *η*(*K* − 1, *s*)} ~ Dirichlet(1,…, 1). Thus, we have 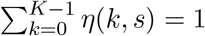 for each sample *s*.
3. Compute the subclonal frequencies *ϕ*(*k, s*) for each subclone *k* in each tumor sample *s* using the tree structure and *η*(*k, s*) values. We have

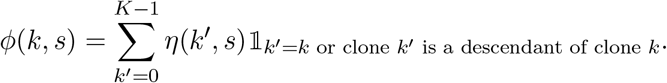
4. Assign the *J* SSMs to subclones. To ensure every subclone has at least one SSM, set the subclones of the first *K* − 1 SSMs λ_s_(0), λ_s_(1),…, λ_s_(*K* − 1) to 1,2,…, *K* − 1. To assign the remaining *J* − *K* + 1 SSMs, sample subclonal weights from the unit Dirichlet, then sample assignments from the categorical distribution using these weights.
5. Segment the genome into I segments by sampling from Dirichlet(1,…, 1).
6. Generate *L* CNAs by assigning each event *l* to a subclone λ_c_(*l*) ∈ {1, 2,…, *K* − 1}, segment *σ*_c_(*l*) ∈ {0, 1,…, *I* − 1}, and phase *π*_c_(*l*) ∈ {*A, B*}. Each assignment is sampled from Dirichlet(5,…, 5) with the appropriate number of dimensions. Subsequently, a direction *d*(*l*) ∈ {gain, loss} is sampled for every doublet (*σ*_c_(l),*π*_c_(*l*)), such that all CNAs with the same segment *i* and the same phase *α* have the same direction. Moreover, deletions are permitted only once on a given tree branch for a given segment and phase.
7. If the direction *d*(*l*) = gain, then the allele gain *g*(*l*) is sampled such that *g*(*l*) ~ ceil(Exponential(λ = 1.5)).
8. If the direction *d*(*l*) = loss, then the allele loss must necessarily be 1, since two CNA events may never have the same segment and phase with opposite directions. This implies that at most one allele can ever be lost.
9. Sample the timing and phase for each SSM *j*. SSM phasing *π*_s_(*j*) ∈ {*A, B*} is sampled from a Dirichlet, such that *π*_s_(*j*) ~ Dirichlet(5, 5). This phasing is rejected and resampled if the given allele and segment has already been deleted in the SSM’s subclone, either in the subclone itself or an ancestor. Subsequently, the SSM’s timing *t*(*j*) is sampled if a CNA has occurred for the same segment and allele in the SSM’s subclone, with *t*(*j*) ∈ {before CNA, after CNA}, and *t* ~ Dirichlet(5, 5). Simulated data parameters are listed in table S3.

**Table S3.** Simulated data parameters. The parameter name in brackets gives the name of the parameter in the simulation script. For datasets without CNAs, we generated a total of one SSM per subclone. For datasets with CNAs, we generated 200 SSMs for each dataset. Assignment of SSMs was performed randomly so that every subclone had a variable number of SSM, but received at least one. Code used to generate the simulated data is available at https://github.com/morrislab/pearsim.

| Parameter | Description | Value |
| --- | --- | --- |
| $K$ | Number of subclones | 5, 20, 50 |
| $N$ | Number of tissue samples | 1, 2, ..., 20 |
| $T$ | Read depth | 50x |
| $J(M)$ | Number of SSMs | 5, 20, 50, 200 |
| $L(C)$ | Number of CNAs | 10, 20, 40 |
| $I(H)$ | Number of genomic segments | 10, 20, 40 |

#### S6.2 Preprocessing of TRACERx data

We worked with the TRACERx data provided in Tables S3 and S7 in the Supplementary Appendix 2 of the work of Jamal-Hanjani et al. [41]. Table S3 contains mutation clusters (column *PyClonePhyloCluster)* and their cancer cellular fraction (CCF, column *PyClonePhyloCCF*) computed by PyClone for 100 patients. After different filtering steps, the authors arrive at 91 patients in Table S7. By avoiding evolutionary conflicts posed by the pigeonhole principle [2] and the crossing rule, and by considering copy number errors, the authors discarded some of the clusters to arrive at a set of mostly consistent clusters (column *TreeClusters* in Table S7) they used to build phylogenetic trees with CITUP [34]. Due to erroneous copy number corrections and a high number of clusters, the authors built manual trees for six patients.

We applied the basic version of SubMARine and used the mostly consistent mutation clusters as subclones and their CCF as subclonal frequencies. Because we did not consider CNAs, we excluded the three datasets with erroneous copy number corrections.

## Notes

### Competing Interest Statement

The authors have declared no competing interest.

